# Uncovering the genetic diversity of the malaria parasite antigen MSP2 across Sub-Saharan Africa

**DOI:** 10.1101/2025.07.23.666329

**Authors:** Julia Zerebinski, Ioanna Broumou, Kelvin M. Kimenyi, Eleni Aklilu, Carolina M. Andrade, Hamza Babiker, Philip Bejon, Mariama K. Cherif, Peter D. Crompton, Cláudia Fançony, Fariba Foroogh, José Pedro Gil, Melissa Kapulu, Steven M. Kiwuwa, Margaret A. Kweku, Anne Liljander, Dinora Lopes, Kevin Marsh, Doreen D. Mutemi, Billy Ngasala, Johan Normark, Judy Orikiiriza, Kristina E.M. Persson, Silvia Portugal, Ulf Ribacke, Sodiomon B. Sirima, Klara Sondén, Taís N de Sousa, Boubacar Traore, Muyideen Kolapo Tijani, Nils Eckerdal, Pär Villner, Lynette Isabella Ochola-Oyier, David F. Plaza, Anna Färnert

## Abstract

Genetic diversity in *Plasmodium falciparum* poses a significant challenge to malaria control and elimination. This is particularly important for developing fully efficacious vaccines, which should include valuable blood stage antigens. Several antigen candidates are highly diverse and require further understanding. We surveyed the genetic diversity of the highly polymorphic merozoite surface protein 2 (MSP2) in 2761 *P. falciparum* isolates collected across Sub-Saharan Africa. Using PCR-based genotyping and long-read sequencing, we identified extensive diversity among *msp2* size variants and sequences. Some size variants were more prevalent than others across different geographical regions, transmission intensities, and time points. These variants comprised multiple unique sequences, of which several were geographically and temporally widespread. Our study reveals greater *msp2* sequence diversity than previously known, while also identifying interesting similarities in sequence and gene length across Sub-Saharan Africa. These findings support the further exploration of common *msp2* variants in relation to parasite virulence and vaccine development.

## Introduction

Malaria remains a significant public health concern globally, with an estimated 263 million cases and 597 thousand deaths in 2023 alone [1]. *Plasmodium falciparum* is the most prevalent and deadly species, accounting for 95% of malaria cases and disproportionately affecting Sub-Saharan Africa. The reduction in incidence that was initiated with introduction of artemisinin combination therapies and vector control strategies has stalled, making new approaches critical [1]. *P. falciparum* is a highly complex pathogen; understanding its spatiotemporal dynamics and genetic diversity is essential for improving surveillance, diagnosis and treatment, as well as for developing preventative measures such as vaccines, to reduce morbidity and mortality and ultimately achieve elimination.

The efficacy of the two vaccines RTS,S/AS01 and R21/MatrixM, targeting the pre-erythrocytic stage [2,3], could potentially be improved with the addition of blood stage antigens. Many of the merozoite surface proteins have a role in host cell binding or invasion [4,5] and are thus interesting targets for antimalarial drugs and vaccines [6]. Among them, the abundant, single copy merozoite surface protein 2 (MSP2) represents a highly polymorphic antigen which has been previously discussed as a valuable target for vaccine development [7]. Such polymorphism serves as a molecular fingerprint [8] which helps determine the number and type of co-existing genotypes present in individual *P. falciparum* infections [9,10], and provides information for malaria control and elimination efforts [11,12]. Accordingly, the World Health Organization recommends using *msp2* in combination with *glurp, msp1,* and microsatellite markers, to track infections and assess drug efficacy by distinguishing between recrudescence and reinfection during drug trials [8,13].

Genotyping of *msp2* is widely used to assess the diversity of the *P. falciparum* parasite population, however, studies using *msp2* as a diversity marker commonly focus on single study sites, reporting multiplicity of infection (MOI), or proportions of the two allelic families [14] in relation to transmission [15,16], age, or malaria severity [17,18]. Few studies have surveyed the diversity of the *msp2* locus over a broad temporal [19,20] or geographical level [21,22]. Such a survey is valuable for understanding the molecular epidemiology and transmission patterns of *P. falciparum*, as well as the distribution of antigenic variants and their importance for immunity.

The *msp2* locus encodes three major structural domains within five blocks: The outer conserved termini are called blocks 1 and 5, and blocks 2 and 4 are referred to as dimorphic regions, containing family-specific sequences used to divide variants into two allelic families FC27 and IC [23]. The central block 3 is highly variable comprising both tandem repeats and variable sequences [23]. While this structure is well documented, no clear patterns of geographical distribution have been identified. Genotyping of *msp2* typically involves PCR amplification of dimorphic and polymorphic blocks 2-4, followed by gel electrophoresis to separate PCR products according to fragment size [24]. While this method is simple and accessible, it lacks the resolution to distinguish between fragments of similar lengths or detect minority clones [25]. The adaptation of fragment sizing to capillary electrophoresis (CE) has improved detection of size variation, enabling highly reproducible sizing of PCR amplicons [26,27]. Although sequencing of *msp2* has previously been limited by gene length and repeats, recent technical advances now allow for long-read sequencing of the entire locus [28,29]. Results from several of our own genotyping studies, as well as others’ studies [15,22], suggest that certain fragment sizes are detected more frequently than others [14,21]. Understanding whether these are common in the parasite population over a large geographic area and over time, is needed for the development of MSP2 as a vaccine target.

## Results

The study included PCR-based genotyping of *msp2* from 2761 *P. falciparum* isolates, originating from 34 countries between 1993 and 2021, in Sub-Saharan Africa (**Figure 1A-C**). Samples were collected as part of 19 studies conducted in 12 countries (n=2391), and from travelers diagnosed with *P. falciparum* malaria in Sweden after arriving from Africa (n=370). Samples were collected primarily from areas of moderate transmission based on parasite prevalence (*Pf*PR_2-10_)(**Table S1)**. After binning to account for technical variability of +/-1 base pairs (**Suppl methods**, **Figure S1**), a total of 6111 *msp2* alleles and 1818 (65.8%) polyclonal infections, with multiple *msp2* alleles were detected.

**Figure 1.**
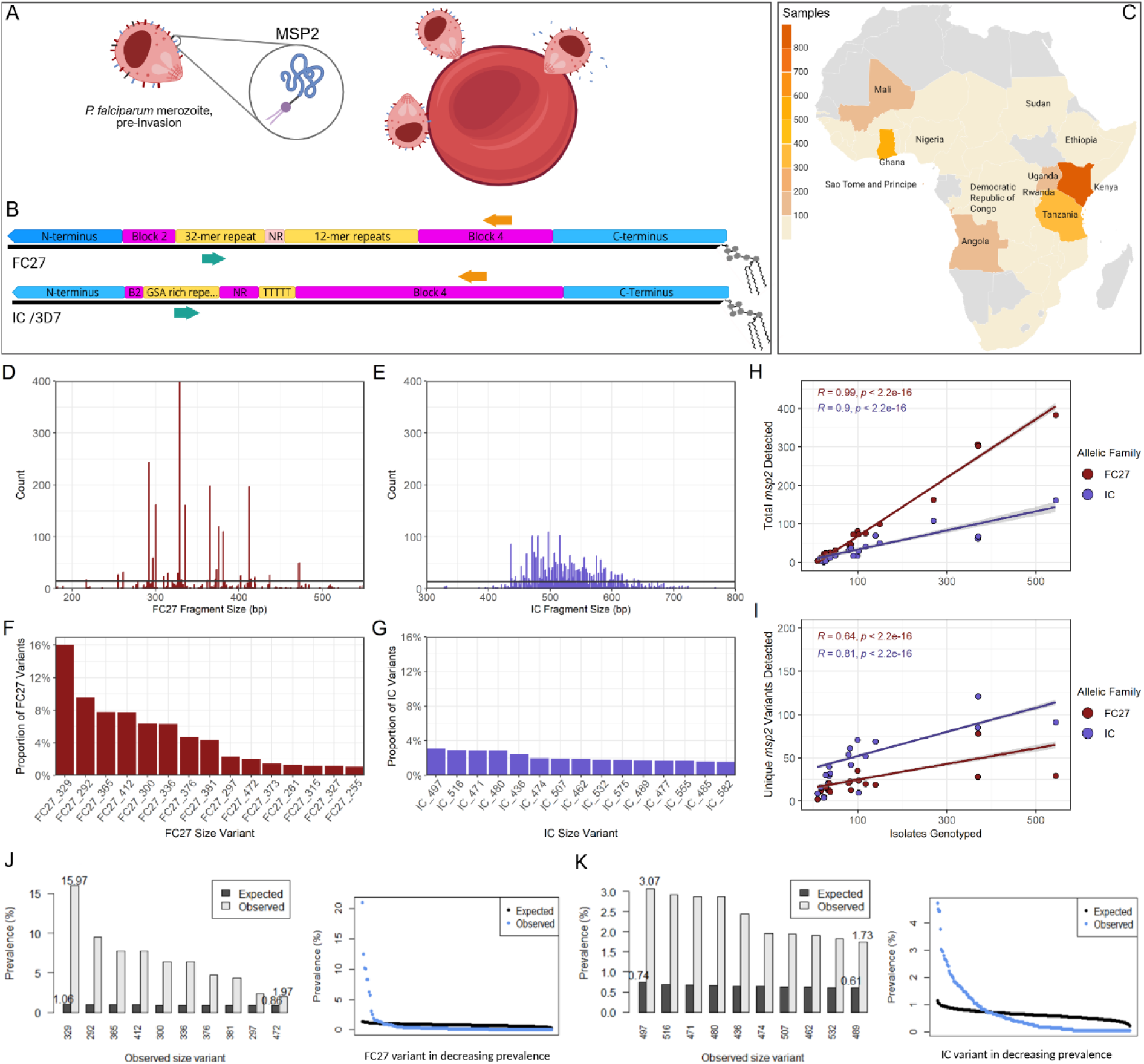
CE genotyping of *msp2* from 2761 *P. falciparum* positive samples collected across 34 countries in sub-Saharan Africa reveals higher prevalence of specific variants. MSP2, the GPI-anchored and second most abundant blood stage surface antigen, is shed from the merozoite during the invasion process (A). The *msp2* locus contains family-specific repeats (yellow), dimorphic blocks 2 and 4 (pink), and conserved termini (blue) (B). Size variants are determined by amplification of the region indicated by forward (turquoise) and reverse primers (orange). Samples were collected from 19 study sites (n=2391) and from travelers returning from sub-Saharan Africa (n=370). Number of *P. falciparum*-positive samples genotyped for *msp2* indicated by color (C). Frequency of *msp2* size variants detected by nested PCR and fragment sizing by capillary electrophoresis, for the FC27 (D) and IC family (E). Grey lines indicate expected frequencies with a population having random distribution of size fragments. Prevalence of the top 15 most frequently detected FC27 (F) and IC (G) variants, in decreasing order of prevalence. The number of total *msp2* size variants detected with increasing sampling effort fitted to a linear model with Pearson correlation (H). The number of unique *msp2* size variants with increasing sampling effort, fitted to a linear model with Pearson correlation (I). The observed prevalence of the 10 most frequent FC27 size variants and all detected FC27 size variants, compared with expected prevalence assuming random chance (Cramér von Mises) (J). The observed prevalence of the 10 most frequent IC size variants and all detected IC size variants, compared with expected prevalence assuming random chance (K).

### Some *msp2* size variants circulate at higher frequency across Sub-Saharan Africa

We identified 412 distinct size variants, including 166 FC27 and 246 IC variants. Distinct peaks in frequency of detection were found for alleles from both families (**Figure 1D, E**). The ten most prevalent FC27 size variants comprised between 3-16% of all FC27 variants (**Figure 1F**), with a cumulative frequency of 67%. The ten most prevalent IC size variants comprised between 2.5-3.5% of all detected IC variants (**Figure 1G**), with a cumulative frequency of 23%. This contrasted with the cumulative prevalence of variants detected once or twice (FC27 n=76, IC n=87), adding up to approximately 3% of variants in each family. The observed distributions of all variants were significantly different from the uniform expected distributions as determined by Cramér von Mises test (FC27 and IC p<0.0001), with expected frequencies of 15 for each unique FC27 variant and 14 for each unique IC variant (grey lines) (**Figure 1D and 1E**).Observed prevalences for the 10 most common FC27 and IC variants were higher than expected (**Figure 1J and 1K**), whereas the observed prevalence of remaining variants declined rapidly to below expected when analysed in decreasing order of prevalence. Study site specific analysis showed non-uniform distributions for the cumulative top 10 FC27 and IC variants at multiple sites (**Table S2**). The total number of detected *msp2* variants was proportional to sampling effort for both FC27 (R=0.99, p<0.0001) and IC (R=0.90, p<0.0001) (**Figure 1H**), as was the number of unique *msp2* variants (FC27 R=0.63, p<0.0001; IC R=0.79, p<0.0001) (**Figure 1I**). Most of the prevalent size variants identified post-binning were also identified at a high proportion pre-binning (**Figure S2A and S2B**).

### Prevalent *msp2* size variants display a wide geographical distribution across Africa

We observed distinct peaks in the frequencies of FC27 (**Figure 2A**) and IC variants (**Figure 2B**) within the respective study sites. By defining unique *msp2* size variants as “species”, we could apply ecological measures of diversity to evaluate similarities between sites. The estimated Sørensen multiple site dissimilarity was 0.23, indicating a high degree of similarity, or overlapping *msp2* variants, between sites. Pairwise Sørensen dissimilarity indices show higher dissimilarity between some sites with higher transmission, but no clear geographical clustering (**Figure 2C**). Alpha diversity values, adjusted for sample size, also showed heterogeneity in the number of unique size variants found at each site, which did not strictly follow transmission intensity (**Figure 2C)**. An accumulation curve modelling unique *msp2* variants with sampling effort begins to plateau, suggesting we have not detected all possible *msp2* size variants (**Figure S2C**). Rarefaction curves for larger datasets, reflect the same plateau tendency but indicate that additional sampling is needed to detect all possible variants (**Figure S2D**). With 412 unique *msp2* size variants, we have detected between 49% and 92% of the estimated total possible species, that is, 486.6 ± 38.48 by bootstrapping, or 763.63 ± 75.85 by Chao 2 measures.

**Figure 2.**
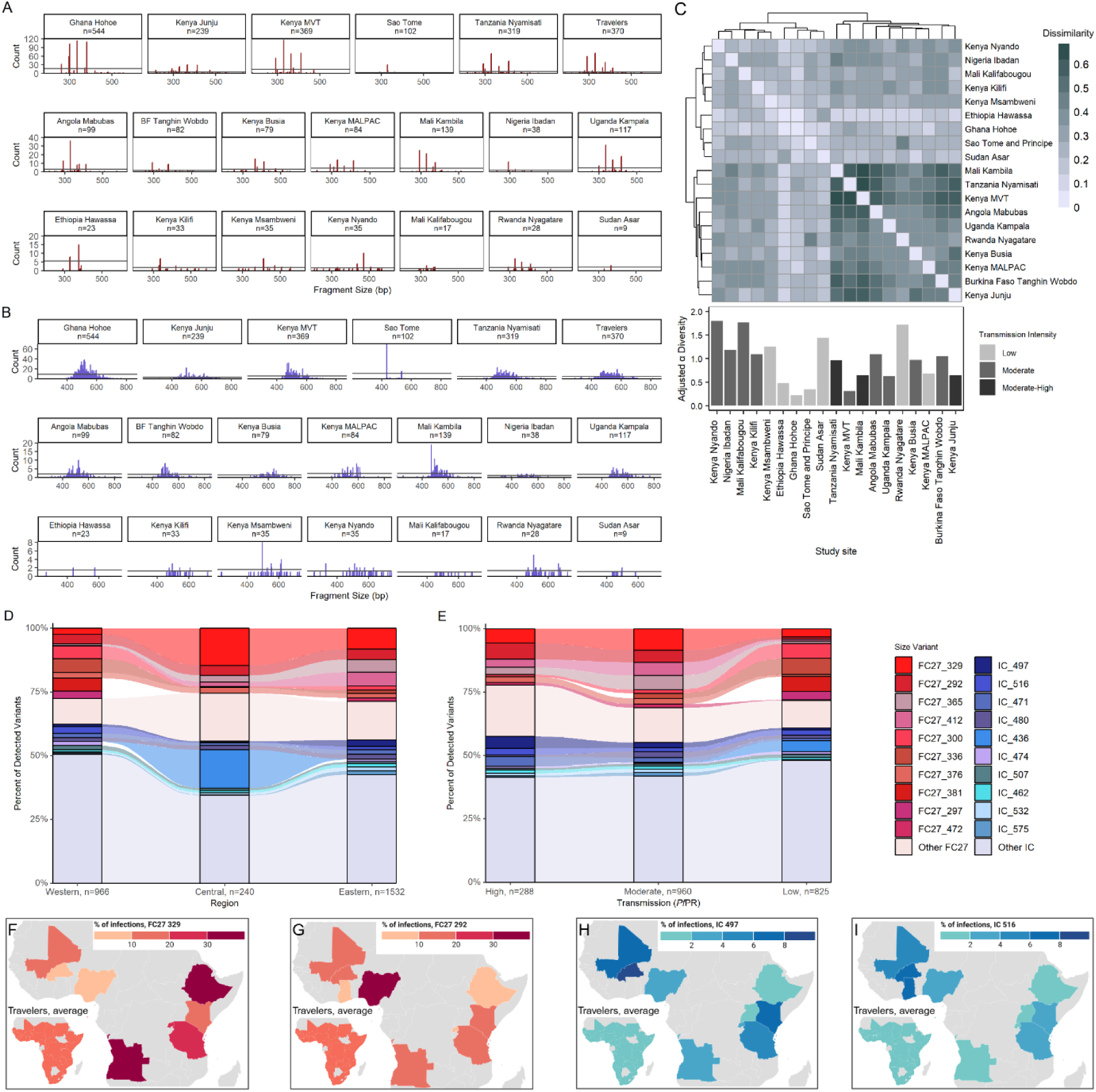
Prevalent *msp2* size variants display a wide geographical distribution across SubSaharan Africa. The detection frequency of size variants, in base pair size, within each study site, shown by histogram for FC27 (**A**) and IC (**B**) variants. The number of genotyped *P. falciparum* isolates at each site is indicated in facet labels; line indicates expected frequencies with a population having uniform distribution. Sørensen beta diversity indices where darker color indicates higher dissimilarity between two sites, and alpha diversity adjusted for sample size shown for each site (**C**). Alluvial plot shows the prevalence of the top 10 most common FC27 and IC size variants as proportion of detected *msp2* variants, within geographical regions (**D**). Alluvial plot shows prevalence of the top 10 FC27 and IC size variants, as proportion of all detected *msp2* size variants, by transmission intensity for study sites not including Travelers, whose exact location of travel was unknown (**E**). Color legend for size variants for both (**D,E**). Transmission intensity was estimated using parasite prevalence data from the Malaria Atlas Project (≥35% *Pf*PR_2-10_ correlates with high transmission, 10-35% *Pf*PR_2-10_ moderate, <10% *Pf*PR_2-10_ low and very low transmission). Prevalence of FC27 329 (**F**), FC27 292 (**G**), IC 497 (**H**) and IC 516 (**I**) in *P. falciparum* positive infections, indicated for each study site and average across all returning Travelers.

The proportion of FC27 and IC variants in the population was highly similar between African regions, although proportions of common variants fluctuated between Western, Central, and Eastern Africa regions (**Figure 2D**). The greater proportions of IC_436 and FC27_329 in Western Africa were mainly due to isolates from São Tomé and Príncipe, analysed further below. The common size variants maintained similar distributions across different transmission intensities (**Figure 2E**). With higher dissimilarity values between moderate and high transmission sites (**Figure 2C**), this suggests the dissimilarity is contributed by variants other than the common ones. When further stratifying into study sites, we observed that several sites had high proportions of samples harboring FC27_329 (**Figure 2F**) or FC27_292 (**Figure 2G**). FC27_329 was detected in a higher proportion of samples than FC27_412 and FC27_300 (p<0.01), and FC27_472 (p<0.05) (**Figure S2E**). Several study sites had around 8% of the population harboring IC_497 (**Figure 2H**) or IC_516 (**Figure 2I**), and overall IC variants were detected at consistent proportions in the population (**Figure S2F**). The proportion of detected variants at each site consisting of the two allelic families was comparable (**Figure S2G**) and the mean MOI by *msp2* was similar across transmission intensities (Kruskal-Wallis p=0.92) (**Figure S2H**).

### Extensive diversity of *msp2* sequences identified by long read sequencing and *in silico* genotyping of *msp2* in 751 isolates

A subset of 1317 samples were subjected to circular consensus sequencing (CCS) followed by genotyping *in silico* using a published and accessible pipeline [28]. Size variants were successfully called from 751 samples (n=1760 sequences) (**Table S3**). After removing artifacts, determined by mapping PCR primers and barcode sequences (**Figure S3A and B**), we retained 1440 sequences from 721 samples. Samples from Kenya Junju (n=60) which exhibited common size variants by CE, were analyzed separately from the remaining 1306 sequences.

Approximately 500 sequences accounted for a size variant within 3 base pairs of the CE call for the same sample (**Figure S3C**). Between 15% and 80% of samples at any given study site had a matching CE and CCS call (**Figure S3D**). We detected 226 distinct FC27 and 463 distinct IC sequences. Until May 2025, 197 full-length *msp2* sequences had been deposited in GenBank, 19 of which were an exact match for 5 sequences in this study: 1 FC27_328, 2 different FC27_292, 1 IC_542, and 1 IC_557; This study detected 684 new, unique, full-length *msp2* sequences. The number of detected full-length *msp2* sequences increased proportionately with sampling effort (FC27 R=0.85, p<0.001; IC R=0.94, p<0.0001) (**Figure 3A**). This was also true for unique, full-length *msp2* sequences (FC27 R=0.79, p=0.0012; IC R=0.9, p=0.0002) (**Figure 3A**).

**Figure 3.**
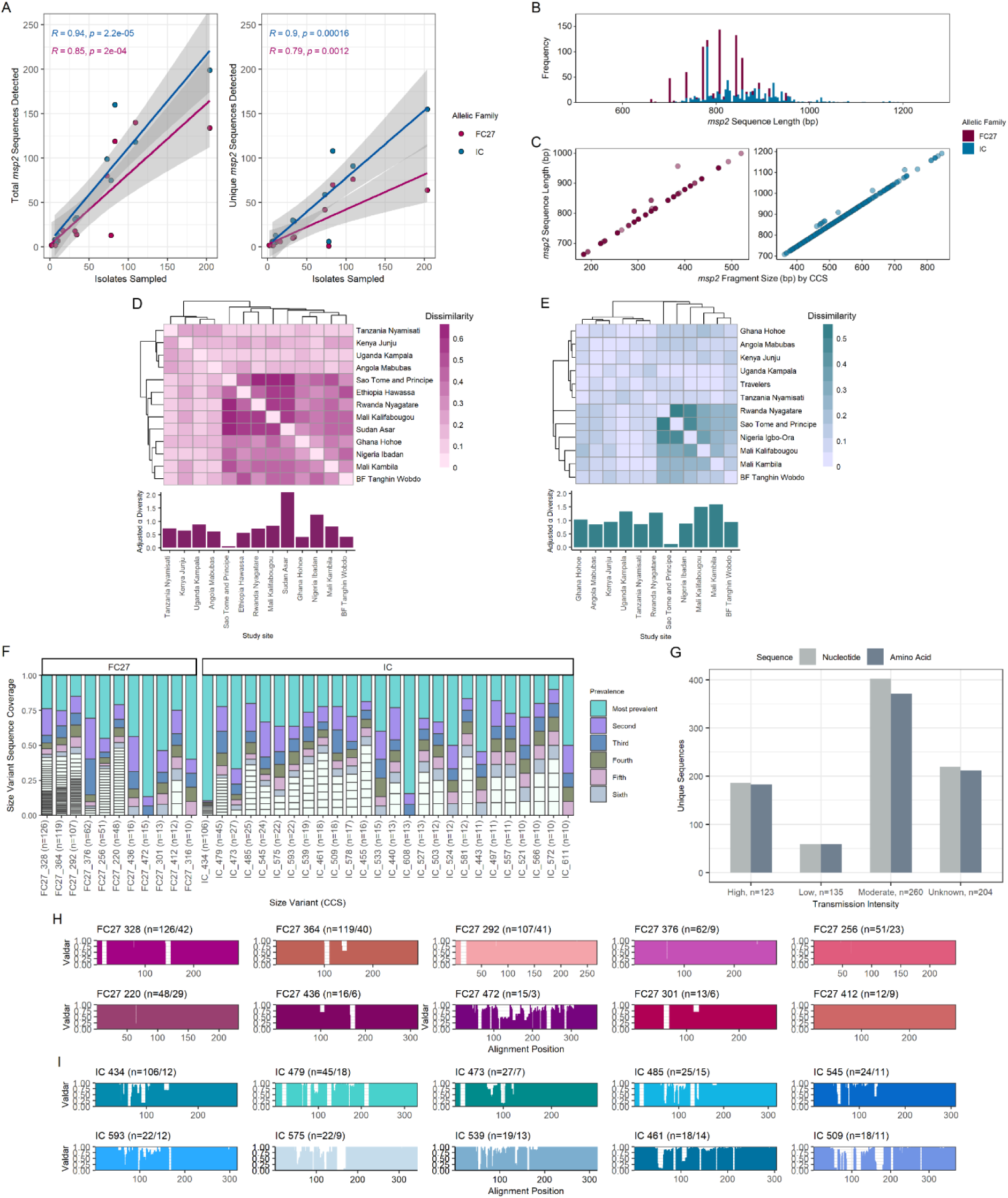
Diversity of *msp2* nucleotide sequences and amino acid sequences coding for size variants. The *msp2* gene was amplified and sequenced. Total detected *msp2* nucleotide sequences and unique *msp2* nucleotide sequences plotted against sequencing effort with linear regression model and Pearson coefficients for both FC27 and IC sequences (**A**), dots indicate study sites. Histogram plotting frequency of detected nucleotide sequence lengths (**B**). Nucleotide sequence length plotted against *msp2* fragment size in base pairs; dots represent single sequences (**C**). Sørensen beta diversity matrices plot dissimilarity in distinct FC27 sequences (**D**) and IC sequences (**E**) between study sites; Alpha diversity based on distinct sequences, adjusted for sample size below (**D**, **E**). Sequence coverage of unique nucleotide sequences out of detected nucleotide sequences for each called size variants detected at least 10 times by CCS-based genotyping (**F**). Number of distinct nucleotide sequences and distinct, predicted amino acid sequences detected, grouped by transmission intensity (**G**). Transmission based on parasite prevalence from the Malaria Atlas Project: ≥35% *Pf*PR_2-10_ high transmission, 10-35% *Pf*PR_2-10_ moderate transmission, <10% *Pf*PR_2-10_ low and very low transmission (n, number of samples in transmission group). Valdar consensus scores for aligned, unique predicted amino acid sequences for the 10 most prevalent CCS-determined FC27 size variants (**H**) and IC size variants (**I**). Total detected and number of unique sequences are indicated for each sub-panel.

We observed distinct peaks in the frequency of CCS-determined fragment sizes (**Figure S4A**). A rarefaction curve indicated that we have not sampled enough to detect all possible sequences (**Figure S4B**). CCS-based genotyping detected similar prevalent *msp2* size variants as PCR-based genotyping (**Figure S4C and S4D**), although fragment lengths differed by 1 or 2 base pairs. For example, FC27_329 (CE) was detected as 328 bp. Additional *in silico* genotyping using the set of primers employed by Mwingira *et al* [21] confirmed the higher prevalence of certain variants in the population (**Figure S4E-G**), however, different fragment lengths were identified due to a different target region.

Full-length *msp2* sequences ranged in length from 663 to 1191 base pairs (**Figure 3B**), and increased proportionately with increasing fragment size (**Figure 3C**). Sequences (dots) above the trend line indicated insertions in the 5’ or the 3’ flank of the size fragment region (**Figure S5A and S5B**). Insertions on the 5’ flank were determined to be 13 or 15 nucleotide inserts separating tandem sites for the nested PCR *msp2* forward primer (**Figure S5C and S5D**) while 3’ inserts were contributed by other indels. Diversity measures were used to describe sequence diversity, where “species” identified unique full-length *msp2* sequences. Sørensen β diversity analysis determined that *msp2* sequence overlap was not clearly dependent on geography transmission intensity for FC27 (**Figure 3D**) or IC (**Figure 3E**). FC27 sequences had overall higher beta dissimilarity between sites, possibly because of the higher number of FC27 sequences detected.

Although multiple distinct sequences were observed for each size variant, we observed that several unique nucleotide sequences were more frequently detected for each size variant (**Figure 3F**). The most prevalent, unique nucleotide sequences coding for FC27_328, FC27_364, and FC27_292 (**Figure S5E-G**) differed from each other in 2 to 60 base pairs (92-99% identity). The dominant sequences for IC_479, IC_485, and IC_473 differed in 1 to 176 base pairs (84-99% identity) (**Figure S5H-J**). Expected heterozygosity values for FC27 size fragment sequences tended to be slightly higher than for full FC27 nucleotide sequences, indicating more SNPs and indels outside than within the size fragment region (**Table S4**). On the other hand, expected heterozygosity values for IC size fragment and full length sequences were more similar to each other, reflecting high diversity in the size fragment region (**Table S4**).

The number of unique, predicted amino acid sequences was as high as for unique nucleotide sequences, demonstrating high contribution of non-synonymous mutations to the generation of diversity in the *msp2* locus (**Figure 3G**). Non-synonymous mutations were common in FC27 sequences (**Figures S6A-C**) while IC family sequences had variable block 3 lengths as well as non-synonymous mutations (**Figures S6D-F**). FC27 family amino acid sequences had varying 32-mer or 12-mer repeats, as expected (**Figure S6G**). IC amino acid sequences demonstrated varying lengths and sequences of the GSA-rich region as well as the number of tandem “T” repeats within a single size variant (**Figure S6H**). The distance between the 3’ end of IC block 3 to the IC reverse primer binding site was also variable within single size variants, although this was most commonly 80 amino acids (**Figure S6H**). We predicted 725 unique protein sequences from 741 unique nucleotide sequences, demonstrating that extensive *msp2* sequence diversity translates significantly into amino acid sequence diversity (**Figure 3G**). To visualize amino acid sequence mutations, we aligned unique predicted protein sequences of commonly detected size variants by CCS and calculated their Valdar conservation scores. The plots show higher conservation in amino acid sequence within FC27 (**Figure 3H)** than IC size variants (**Figure 3I)**, even with the overall higher detection rate of FC27 type sequences. For example, FC27_376, FC27_256, FC27_220, and FC27_412 show almost 100% conservation at each amino acid residue of their alignments, which was not observed for IC amino acid sequences.

### *Msp2* sequences are widely distributed across Sub-Saharan Africa

We evaluated the relatedness of all full length *msp2* sequences using maximum likelihood phylogenetic trees. Neither FC27 (**Figure 4A**) nor IC sequences (**Figure 4B**) clustered geographically, whether by country or by African region. Unique full length sequences for common *msp2* variants were geographically widely spread, especially the most prevalent FC27 sequences (as identified in **Figure 3F**) at the top of each facet (**Figure 4C**). This pattern was also evident in unique IC sequences (**Figure 4D**), although they were detected in fewer samples and thus appeared to be less geographically widespread than FC27. To ensure that the unstructured phylogenies were not due to conserved regions of *msp2*, we trimmed all full-length sequences to include only the size fragment region. We then performed a principal component analysis (PCA) on size fragment sequences. The allelic families clustered together but showed no geographical clustering (**Figure 4E**). FC27 size fragment sequences (**Figure 4F**) and IC size fragment sequences (**Figure 4G**) clustered mainly by *msp2* size variant (size fragment length). A PCA of size fragment sequences from FC27_328 (**Figure 4H**), FC27_292 (**Figure 4I**), IC_479 (**Figure 4J**), and IC_473 (**Figure 4K**) alone, showed no geographical clustering.

**Figure 4.**
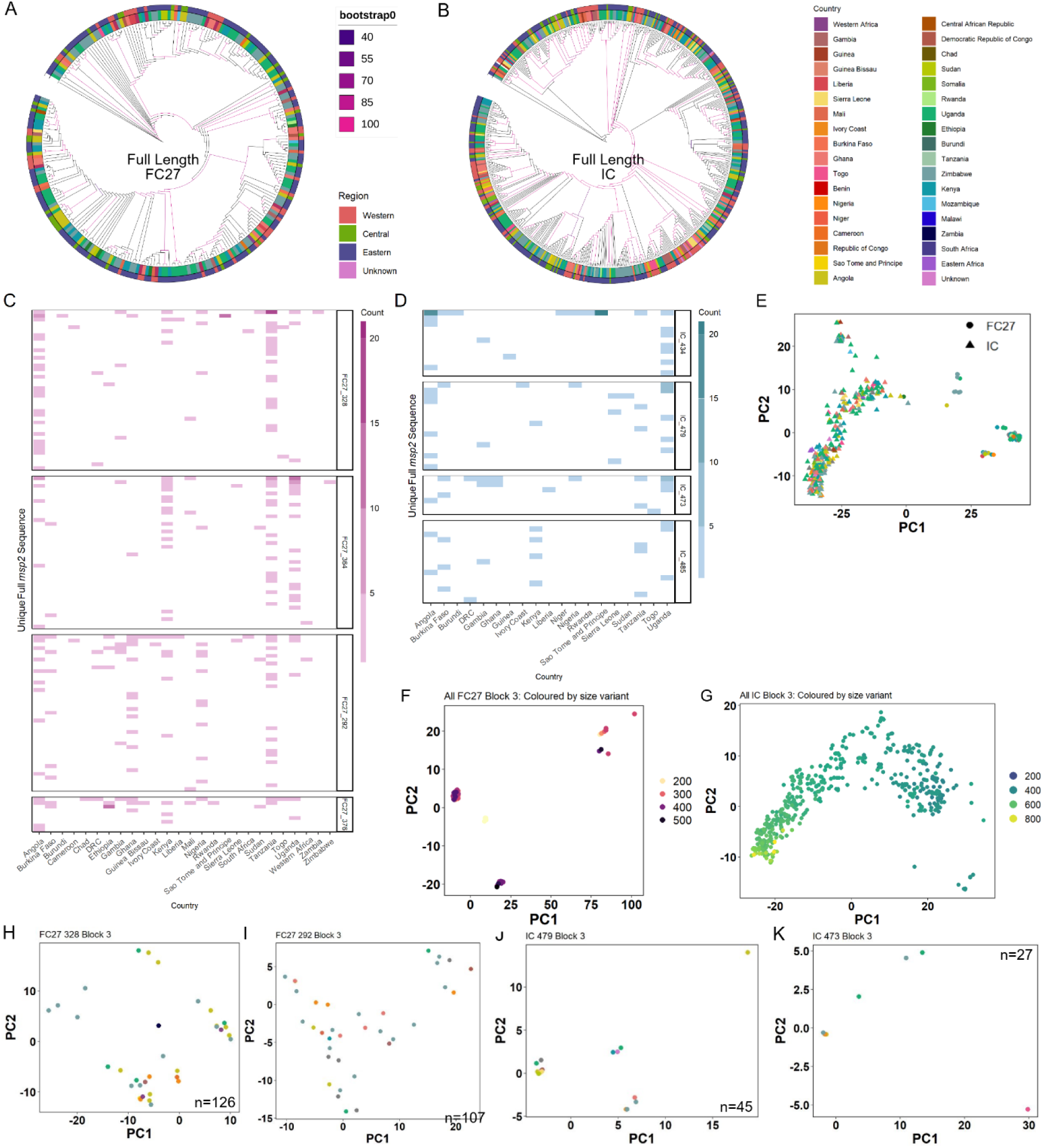
*Msp2* sequences are geographically widespread across SubSaharan Africa. Phylogenetic trees were constructed for all detected full length *msp2* nucleotide sequences detected for FC27 (**A**) and IC (**B**), using *P. billcollinsi* G01 *msp2* as the root. African region (outer ring) and country of isolate (inner ring) indicated by color legends. Bootstrap support indicated by colored branches. Distinct sequences detected for frequently detected (CCS) FC27 (C) and IC (D) size variants were plotted by country of isolate; Frequency indicated by strength of color. Full length nucleotide sequences were trimmed to contain only the region targeted by fragment size oligos. Principal component analysis (PCA) was performed on all size fragment sequences (**E**), allelic family indicated by shape. All FC27 size fragment sequences (**F**) and all IC size fragment sequences (**G**) were plotted by PCA and colored by size variant. Size fragment sequences corresponding to FC27 328 (**H**) and FC27 292 (**I**) size variants were separately plotted by PCA. Size fragment sequences corresponding to IC 479 (**J**) and IC 473 (**K**) were also plotted by PCA and colored according to country, as indicated by the color legend. Number of size fragment sequences indicated.

### Maintenance of *msp2* size and sequence variants over time within a single population

In one site, Nyamisati, a village on the coast of Tanzania (**Figure 5A**), repeated sampling during a high transmission period (1999) and a low transmission period (2016) [30] revealed a stable distribution of *msp2* size variants by CE over time (**Figure 5B**). Common variants remained consistent over the two survey years, with a slight overall increase in IC variants. Several variants, including FC27_412, FC27_329, FC27_292, and IC_509, were detected at a relatively high proportion of tested individuals (15-25%) in both survey years (**Figure 5C**). We then examined whether *msp2* sequences were conserved over time. Several sequences, including FC27_292, FC27_364, IC_497, and IC_485, persisted over the two periods (**Figure 5D**). A maximum likelihood phylogenetic tree showed no clear temporal clustering for full-length *msp2* sequences in the village (**Figure 5E**), but grouped sequences by fragment size (**Figure 5F and 5G**). PCA of *mps2* size fragment sequences from Nyamisati residents and travelers to Tanzania revealed no clustering for FC27_328 (**Figure 5H**) or for FC27_292 (**Figure 5I**), indicating no significant differences between the parasite *msp2* genotyped from the two groups.

**Figure 5.**
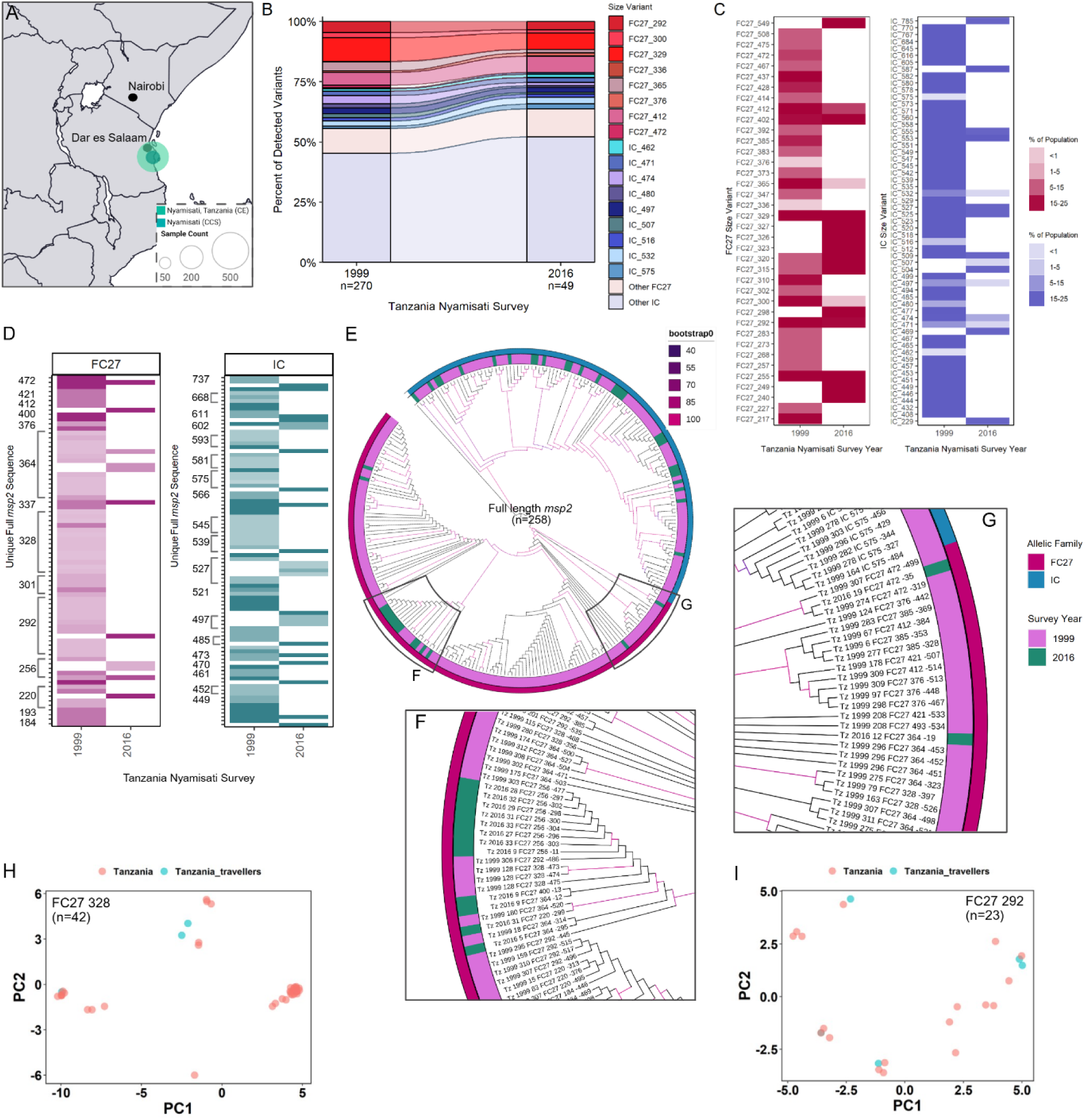
Distribution of *msp2* size and sequence variants over time in a single site in Tanzania. Geographical location of Tanzania Nyamisati, with number of samples genotyped by CE and CCS indicated by size of colored dot (**A**). Distribution of *msp2* size variants (CE) in Nyamisati 1999 and 2016, expressed as a proportion of all detected *msp2* variants (**B**). Prevalence of *msp2* variants in the population, as percentage of individuals harboring each variant in 1999 and 2016 surveys (**C**). Unique, full length *msp2* sequences detected in Tanzania Nyamisati were plotted across the two survey years, where coverage indicates the proportion of each unique sequence out of total sequences identified for single size variant (CCS)(**D**). A phylogenetic tree was constructed using all full length *msp2* sequences isolated from Nyamisati (**E**). Inset panels show clustering by size variant rather than by survey year (**F and G**). Colors indicate each survey year (inner ring) and *msp2* allelic family (outer). Branch color indicates bootstrap support. Size fragment sequences from FC27 328 (**H**) and FC27 292 (**I**) were plotted by PCA, where colored dots indicate sequences isolated from Tanzania Nyamisati samples (coral) or travelers returning from Tanzania (teal). Dots are jittered to avoid overlap of identical sequences.

### Low *msp2* diversity in an isolated, low transmission setting

To illustrate an isolated population with low transmission, we examined São Tomé and Príncipe in more detail (**Figure 6A**). Using CE, we detected 187 *msp2* alleles from the 102 samples, of which 34 were distinct. A minority of variants (IC_436, FC27_329, IC_535, and FC27_331) were highly prevalent in both the total *msp2* alleles detected (**Figure 6B**) and in *P. falciparum* infections (**Figure 6C**). More than 30% of individuals (32 of 103) carried both IC_436 and FC27_329. We successfully sequenced *msp2* from 73 samples, retrieving 101 full-length sequences in total and 7 unique full-length sequences, demonstrating markedly low sequence diversity within the detected variants (**Figure 6D**). Most of the sampled individuals carried IC_434 by CE and by CCS; 5 individuals (6.8%) carried both IC_434 and FC27_328 by CCS. Although a single IC_434 *msp2* sequence was found circulating in the São Tomé and Príncipe samples, it was not unique to São Tomé and Príncipe: the same unique sequence was detected in samples from multiple countries, resulting in a geographically unstructured phylogenetic tree (**Figure 6E**).

**Figure 6.**
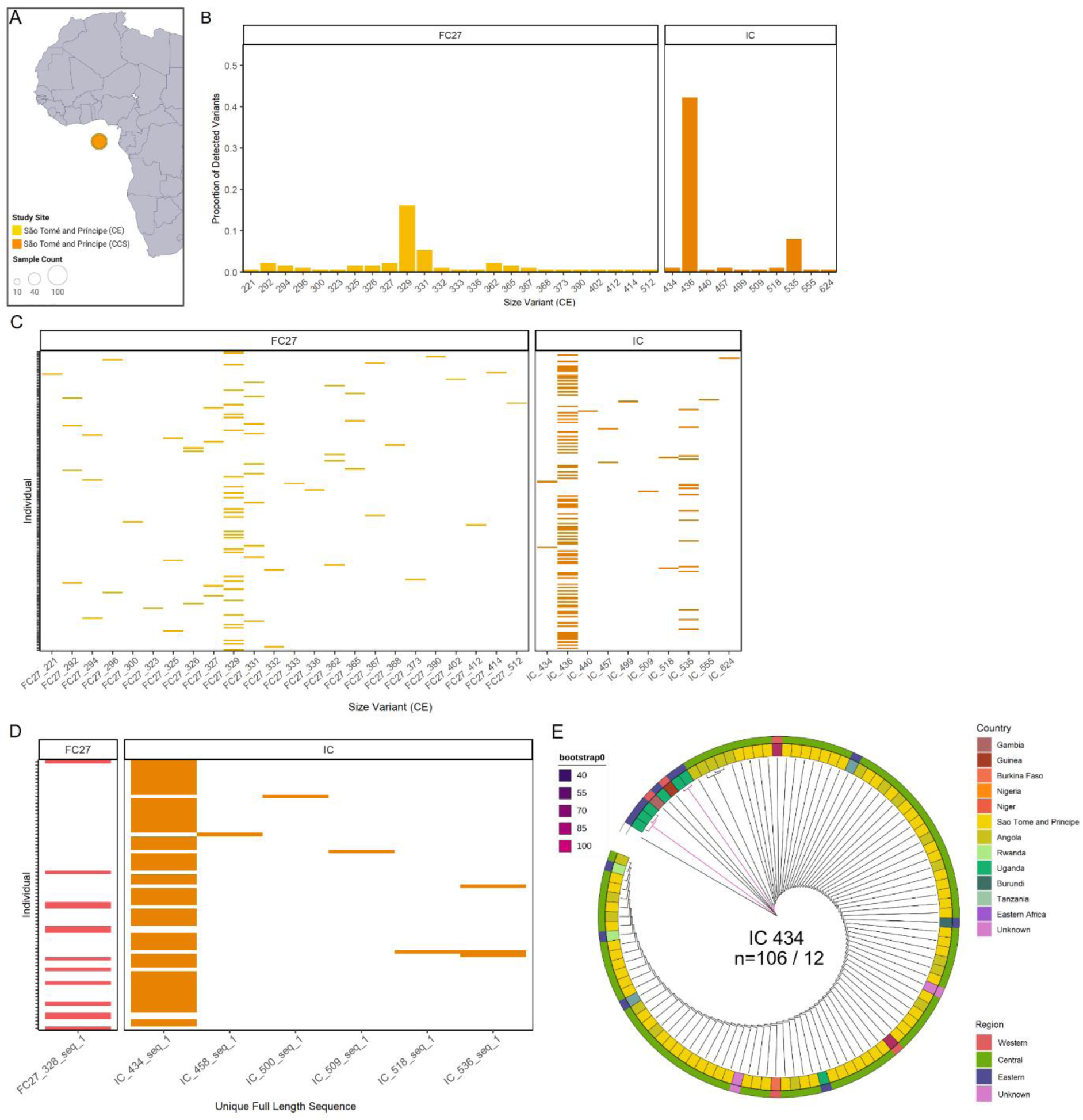
Distribution of *msp2* variants in the isolated, low transmission site of São Tomé and Príncipe. The most isolated study site, Sao Tome and Príncipe, is an island with low transmission; Number of samples genotyped by CE and CCS indicated by size of colored dot (**A**). Observed *msp2* size variants by CE and their prevalence as proportion of all detected *msp2* variants in São Tomé and Príncipe (**B**). Observed *msp2* size variants by CE detected in each individual sampled (**C**). Unique sequences detected within each sample (**D**). All full length *msp2* nucleotide sequences called IC_434 by *in silico* genotyping were aligned and a phylogenetic tree was constructed with root at *P. billcollinsi* G01 *msp2* (**E**). Colors indicate country of isolate origin (inner ring) and African region (outer ring). Branch colors indicate bootstrap support. Total number of sequences and number of unique sequences indicated.

### *Msp2* size variants prevalent across Kenyan study sites

Samples from Kenya (n=874) accounted for 31.7% of the study population, allowing a regional comparative analysis. Junju (n=608), Kilifi (n=117), and Msambweni (n=35) are located on the southern coast of Kenya while Nyando (n=35) and Busia (n=79) are located to the west; Travelers who recorded Kenya as country of visit (n=29) were also included (**Figure 7A**). FC27 *msp2* size variants (CE) were widely distributed among the sites and in travelers, with many sites showing 15-20% coverage (**Figure 7B**). FC27_365 was detected in all sites; FC27_412, FC27_329, and FC27_292 were detected in 6 of 8 sites including travelers (**Figure 7B inset**). IC_532, IC_516, and IC_497 were detected in at least 6 of the 8 sites; and IC_529 and IC_462 were detected in 7 of the 8 sites (**Figure 7C**). The study in Kenya Junju assessed children through yearly cross-sectional surveys from 2007 to 2018 [20]. To further evaluate *msp2* dynamics in this population over time, we included repeated sampling of the individuals analyzed in this study. With all included samples (n=472), we observed an increase in the commonly detected CE size variants, for example FC27_329, FC27_365, and FC27_412, between 2007 and 2018 (**Figure 7D**). We observed maintenance of the common IC variants over time, although not all 10 were detected in Kenya Junju. The overall proportion of FC27 and IC alleles fluctuated over the survey years (**Figure 7D**). The same fluctuation in allelic family distribution, and increasing prevalence of common variants, was detected without repeated sampling of the same study participants (**Figure S6I**). A subset of samples from Kenya Junju, with commonly detected size variants by CE, were selected for sequencing. Similar to the dynamics within Tanzania Nyamisati, unique *msp2* sequences were maintained in the population over several surveys (**Figure 7E**). A maximum likelihood phylogenetic tree did not show clustering of full length *msp2* sequences by survey year (**Figure 7F**). A PCA comparing FC27_292 size fragment sequences from Kenya Junju those from travelers, appears to cluster the sequences (**Figure S6J**), however, the number of sequences was small. The absence of clustering and the detection of unique *msp2* sequences across Kenya and over time in Junju, demonstrate the maintenance of common variants in the parasite population.

**Figure 7.**
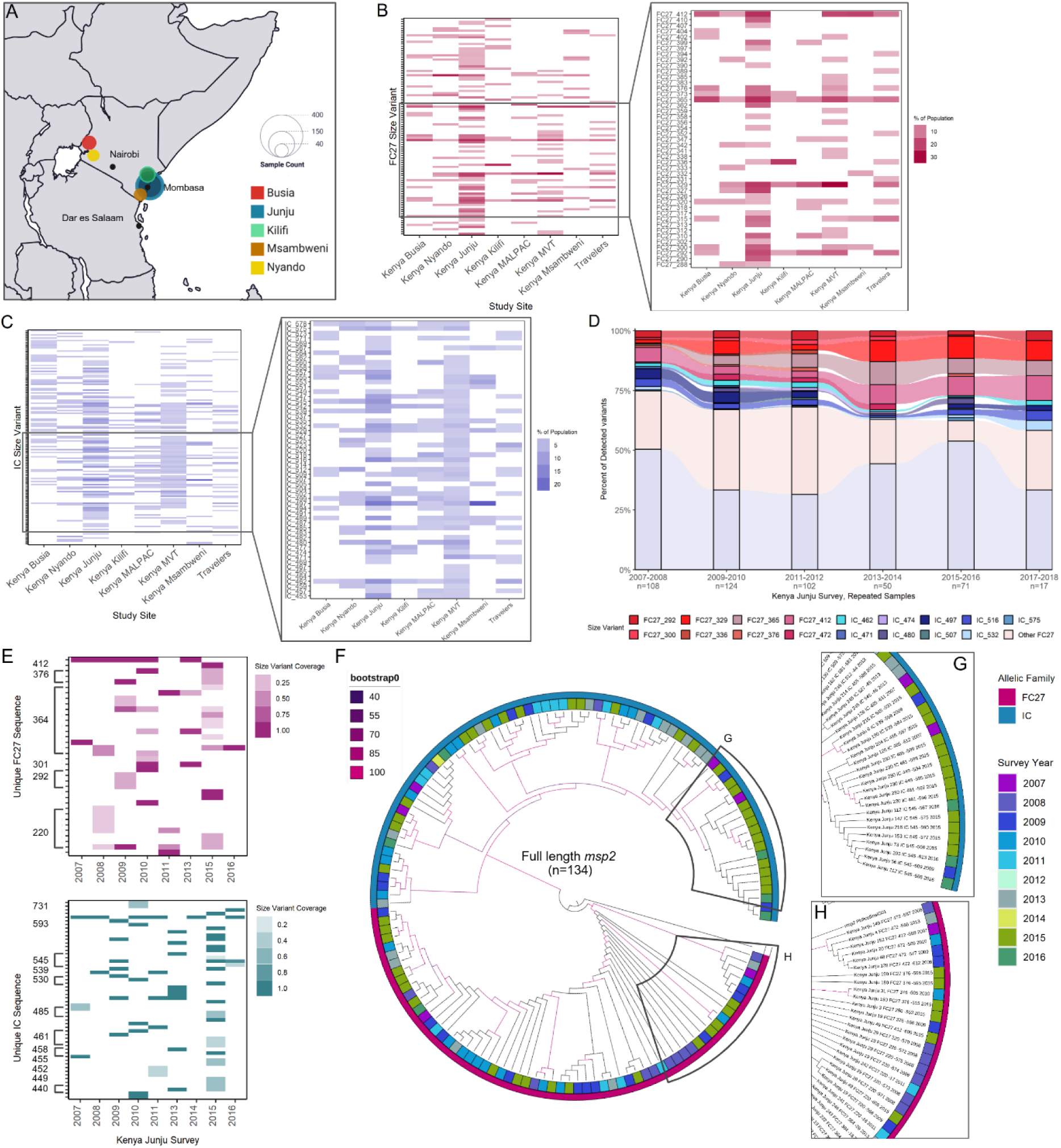
Distribution of *msp2* size variants throughout Kenyan study sites. Geographical location of Kenyan study sites (Travelers not indicated) where number of samples is denoted by size of colored dots (**A**). Proportion of samples (*P. falciparum* infections) per Kenyan study site and in Travelers, harboring each FC27 size variant (CE) **(B)**, with prevalent variants shown in inset. Proportion of samples per study site and in Travelers, harboring each IC size variant (CE) **(C)**, with prevalent variants highlighted (inset). Distribution of *msp2* size variants (CE) in the parasite population of Kenya Junju over grouped survey years (**D**). Individuals who had repeated sampling at Kenya Junju (n=472) were also included for the alluvial plot. Unique, full length *msp2* sequences detected in Kenya Junju were plotted across survey years (**E**), with corresponding size variants (CCS) indicated on the y axis. Sequences from Kenya Junju, which were a selection of samples harboring common size variants identified by CE,were aligned with MAFFT and a phylogenetic tree was constructed **(F**). Colors indicate survey year (inner ring) and *msp2* allelic family (outer ring). Branch color indicates bootstrap support.

## Discussion

In this study, we performed a comprehensive analysis of the genetic diversity of the *msp2* locus in 2761 *P. falciparum* infections from across Africa, combining genotyping by nested PCR and fragment sizing by capillary electrophoresis and high-fidelity long-read sequencing. We identified the most prevalent *msp2* size variants, which were maintained across geographical areas, temporal groups, and transmission intensities. We reveal the extensive diversity of both nucleotide and protein sequences coding for single size variants, although certain sequences were more prevalent than others, and widespread both geographically and temporally. Finally, despite extensive *msp2* diversity, we demonstrate a pattern of maintenance of certain *msp2* size variants and sequences in the parasite population. We believe these variants may be a target for vaccine development, and as such should be further studied in relation to parasite virulence and human immunity.

From the 412 unique *msp2* size variants, the most prevalent were found across all survey sites. While the frequency of detection for the ten most common variants from each family differed between study sites, they were consistently more prevalent than other variants. Common size variants from both FC27 and IC families were detected at high frequencies (10-30%) across multiple sites. In Tanzania and Kenya, repeated surveys showed that common variants were maintained over time and declining transmission intensity. Within an extensive diversity of full-length *msp2* sequences, certain sequences were found to be more prevalent than others. These were detected across multiple survey sites and countries, and repeated surveys in Tanzania and Kenya, demonstrating a pattern of *msp2* length and sequence maintenance in the parasite population. Importantly, our report of prevalent *msp2* variants is supported by previous studies showing a higher prevalence of certain variants [14,21]. Additionally, a previous study observed the same *msp2* size variants and distribution between the transmission season and the dry season in Mali [31]. A large proportion of infections within many study sites (10-40%) carried FC27_329 or FC27_292 size variants, which could indicate a target for vaccination. Further analysis of the antigens coded by the sequences included here, and the immune responses towards them, will help to increase the understanding of which variants could be included in a multivalent vaccine to provide significant population coverage.

In contrast to the extensive diversity found across Africa, we observed very low *msp2* diversity in São Tomé and Príncipe, a highly isolated island nation characterized by low *P. falciparum* transmission for the last 15 years [32]. Among the detected *msp2* variants, 40% were IC_436, and almost 70% were either IC_436, IC_535, or FC27_329. A large proportion of samples carried IC_436 either alone or in combination with FC27_329 or another variant. While lower diversity might be expected in this kind of setting, we were surprised to see the same genotype, and combination of genotypes, in the entire population. The finding was supported by sequencing, which detected only a single unique full length *msp2* sequence for each of the 7 variants identified by CCS. This indicates that there are very low recombination rates on the island. Possible explanations include recent introduction of the circulating parasites, lower immune pressure resulting from low transmission intensity, or even accessible treatment which might limit parasite recombination and drive the selection of specific parasite populations.

Previous studies reported low diversity for the island nation [33], with suggested importation from West or Central Africa [32]. Similar findings have been shown in Cabo Verde following the re-introduction of parasites [34]. More recently, low diversity in the parasite population was indicated by the dominance of 4-aminoquinoline resistance associated alleles, likely due to selection bottleneck events (Té *et al*, submitted). We were unable to determine whether the dominant *mps2* variants were imported from specific areas, since their sequences were detected in samples from multiple African countries.

Our findings revealed extensive *msp2* sequence diversity in nature, representing a challenge to the use of fragment size-based genotyping. Identical fragment sizes may represent different sequences, resulting in under-estimation of the number of co-existing clones and miss-typing of reinfections. In MSP2-based vaccination trials, breakthrough infections of seemingly homologous strains may, in fact, be variants of homologous length [11] but different sequence. Thus, additional markers to discriminate between reinfection and recrudescent strains remain necessary. Furthermore, a threshold number of SNPs or percent identity should be determined for calling sequences as different variants. Sequencing methods, such as those using PacBio, have historically been less accessible to those performing regular surveillance in low-income, high-malaria-burden countries. However, recent developments in nanopore technology allow sequencing at a lower cost than previously possible [29,35]. Challenges remain for widespread implementation, but this technology is likely the way forward when sequence-level precision is needed. Single polymorphisms in sequences coding for common variants may impact human immune recognition, since single amino acid mutations can affect antibody recognition and affinity [36]. Our identification of an almost 1-to-1 ratio of unique nucleotide sequences to unique amino acid sequences showcases a significant hurdle to immune recognition of merozoites.

The interpretation of population structure and diversity based on *msp2* is not optimal as the locus is under strong immune selection [37]. Nevertheless, we were interested in mapping the diversity on a geographical scale given that MSP2 is associated with immunity and has been tested as a vaccine candidate [38–41]. Sequence diversity is advantageous for immune evasion, helping to distract immune attacks and leading to strain-specific immunity [42,43]. FC27 *msp2* variants with differing numbers of tandem repeats have been shown to induce varying levels of antibody response [44–46]. The most common FC27 size variants in our study (328 bp, 292 bp, and 365 bp by CE) correspond to increasing copy numbers of 12-amino acid tandem repeats.

However, sequence diversity within these variants could influence recognition of both linear and conformational epitopes. Importantly, even with extensive sequence diversity the common size variants persist in the population, highlighting some importance of gene length. Irrespective of the extensive *msp2* sequence diversity reported here, size variants and unique sequences were found in different locations and were maintained over time. Individuals do become immune to clinical malaria after repeated infections, without encountering all possible variants of MSP2 and of other variable antigens. Responses against a combination of variants, such as the most prevalent variants described here, may elicit sufficient protection. Previous work has shown that cross-recognition of different variants is already detected in plasma from travelers, even though the conserved termini are not targeted by IgG antibodies [47]. Further study is necessary to understand the impact of sequence or gene length diversity on immune recognition. This will help us determine which MSP2 alleles could be combined in a multicomponent vaccine.

While structural modulations of recombinant MSP2 have been studied [48,49], little is known about its conformation or role on the parasite surface. High affinity antibodies towards MSP2 and other merozoite antigens are associated with protection [50], but importantly, antibody affinity is impacted by conformational changes in the antigen [51]. MSP2 is a highly abundant surface protein, and is thought to play a role in lipid interactions [48,52,53]. Although allelic dimorphism is not unique to *msp2* [54], its specific role in *P. falciparum* remains unclear. The kinetics of fibril formation in FC27 and IC MSP2 differ [55], due to variation outside the conserved N and C termini [48]. Many intrinsically disordered proteins contain short secondary structure regions flanked by disordered regions, called molecular recognition features [56].

Typically involved in molecular interactions, these regions can induce folding [57] and are prone to aggregation [58], a phenomenon that has been noted for MSP2 [49,59]. The secondary structure resulting from MSP2’s polymorphic block 3 and flanking regions may have such a role in modulating binding, influencing fibril formations, or hiding epitopes [57,58,60,61]. MSP2 also appears to be dispensable [62], suggesting redundancy or a role in immune evasion. Indeed, the highly observed non-synonymous mutations in our data are common to genes subject to strong selective pressure, particularly those implicated in immune evasion [63]. While immune pressure drives the extensive sequence diversity, the persistence of certain size variants suggests that gene length may have an important role. Common variants, like FC27_329, could be more easily expressed and maintained in the population, with mutations influencing immune recognition but not gene length.

There are several limitations to this study. First, we were unable to retrieve all sequenced *msp2* from the *in silico* pipeline due to the noise threshold, based on negative controls. While this may have removed true reads, it also ensures that all output sequences are true reads. Secondly, our statistical analysis was based on the hypothesis that any *msp2* variant genotyped from a randomly distributed population would have highly similar prevalence. This assumption does not account for the variable lengths of blocks 2 and 4, the additional family-specific repeats, or increased numbers of T’s in the IC block 3. Without knowing the mechanism behind why or how these kinds of repeats are added or deleted from the locus, we could not factor this into the statistical analysis. Finally, different sample materials, equipment, and slightly different PCR conditions between laboratories may have affected our ability to detect and genotype parasite *msp2* with equal accuracy and precision. While it is possible that we missed minority clones with, for example dried blood spot samples, our main observations were not limited to sites with the exact same sampling methods. On the other hand, the major strength of this study is the broad geographical and temporal spread of samples, which we believe makes it representative of *P. falciparum msp2* across Sub-Saharan Africa. The precision and reproducibility of length genotyping by CE, and the novel genotyping method based on high-fidelity long-read sequencing, are also major strengths. While our results confirm the diversity in *msp2* gene length, our study shows for the first time the extensive sequence diversity in single CE-determined size variants. Our results provide a foundation for further research and a better understanding of polymorphic antigens, which are important not only in *P. falciparum* but in other human pathogens as well.

To conclude, we present a comprehensive study of *P. falciparum msp2* genetic diversity across Sub-Saharan Africa. We identified extensive diversity in both *msp2* gene length and sequence. We also demonstrate a pattern of maintenance of the most commonly detected *msp2* size variants and sequences across a broad geographical and temporal scale. We believe that our results lay groundwork for identifying which variants are important for parasite biology and naturally acquired immunity to malaria. Furthermore, our findings suggest features of a polymorphic locus which could be leveraged for vaccine design, and may also be applicable to other polymorphic antigens. Additional studies of common *msp2* variants and sequence diversity in the context of immunity, could pave the way for the further development of MSP2 as a multicomponent vaccine target.

## Methods

### Ethics statement

This study was approved by the Swedish Ethical Review Authority (Dnr 2008-998-31-3; Dnr 2011/1658-31/4 with amendments 2018/1967-32, 2024-01464/2; 2019-05746). All included studies were approved by the relevant national and institutional ethical review boards in the respective countries. Details of ethical approval for published studies can be found in the original studies (**Table S1**). For the following unpublished studies: Sample collection and analysis of São Tomé and Príncipe isolates, was approved by the IHMT-ITQB Ethics Committee (ref. 16.22) and Swedish Ethical Review Authority (ref. 2017/499-32); The study in Burkina Faso (Tanghin Wobdo) was approved by the Comite d’ethique pour la recherche en sante, Burkina Faso (Nr 2020-4-067); All sample data was pseudonymized, without access to information about the donors other than their location, year, age and the type of study.

### Study populations and sample selection

The raw *msp2* CE genotyping data analysed in this study was either previously published, available but not detailed in published studies, or produced for this study. Studies have been described in detail in the original publications (**Table S1).** Travelers included individuals returning to Sweden with acute *P. falciparum* malaria, treated at Karolinska University Hospital (n=324) [64,65], and migrants arriving to Sweden from Africa and testing positive for *P. falciparum* in a screening study (n=46) [66]. Studies from Burkina Faso, Ethiopia Hawassa and São Tomé and Principe, are in preparation or submitted.

Samples were included if they tested positive for *P. falciparum* by microscopy and/or PCR. Genotyping data inclusion was based on CE positivity to ensure detectable reads (**Table S1**). In studies with repeated sampling of the same individuals, only the chronologically first *P. falciparum* positive sample was included. For Kenya Junju, a subanalysis included data from cross-sectional surveys over consecutive years, however, the main analysis included only first positive samples. In studies with drug treatment, only placebo group samples or baseline samples were included. For Tanzania Nyamisati two cross sectional surveys (1999 and 2016) were selected from a longitudinal population study [30,67]. Estimates of transmission intensity were obtained from the Malaria Atlas Project by *P. falciparum* parasite rates in children 2-10 years old (*Pf*PR_2-10_). Village or city coordinates (latitude and longitude) were used to obtain parasite rates at study sites, rather than study prevalence which was not available for all studies. Exact locations of travelers’ visits were not known, so *msp2* data from this group was not included in the analysis of transmission intensity.

## DNA extraction

Original sample material and extraction methods used for the respective studies are summarized in **Table S1**. For most studies, DNA was extracted from venous blood or dried blood spots (DBS) using QIAamp DNA mini kit (Cat No. 51106) according to manufacturer instructions, with the modification of using 100ul of elution buffer to increase DNA concentration. For DBS extraction, 3 punches of 3mm were used except for samples from Ethiopia Hawassa, where 6 punches of 4mm were taken. Some studies used ABI Prism 6100 Nucleic Acid PrepStation (Applied Biosystems) or Chelex extraction (**Table S1**).

### PCR based *msp2* genotyping and fragment sizing by capillary electrophoresis

Genotyping of *msp2* was based on nested PCR followed by fragment sizing by capillary electrophoresis as previously described [26]. Briefly, a primary PCR amplified the entire *msp2* open reading frame, where each 10ul reaction included 2ul 10x Buffer II, 1.6ul 25mM MgCl2, 0.25ul 10mM dNTP mix, 0.5 ul each 10uM forward and reverse primer (Invitrogen), 0.08ul AmpliTaq DNA Polymerase at 5 units/ul, and 3ul template DNA. The PCR conditions included an initial denaturation of 5 minutes at 95℃, 25 cycles of: annealing 2 minutes at 58℃, extension 2 minutes at 72℃, and denaturation 1 minute at 94℃; then annealing for 2 minutes at 58℃ and extension 5 minutes at 72℃. This was followed by two separate, secondary PCR reactions, each targeting the FC27 and IC dimorphic sequences (**Figure 1B**) using fluorescent family-specific primers (Invitrogen), where each 10ul reaction included 2ul 10x Buffer II, 0.8 ul 25mM MgCl2, 0.25ul 10mM dNTP mix, 0.25 ul (FC27) or 0.6ul (IC) each 10uM forward and reverse primer (Invitrogen), 0.08ul (FC27) or 0.2ul (IC) AmpliTaq DNA Polymerase at 5 units/ul, and 1ul amplified DNA from reaction 1. A slight modification was included [27] for samples analysed in-house (10 cohorts) (**Table S1**), consisting of an increase in the temperature of the nested reaction annealing step from 58℃ to 61℃. Detection and sizing of amplified PCR products CE was done using ABI 3730 PRISM® DNA Analyzer and 50cm capillary length, with LIZ 1200 size standard. GeneMapper version 5.0 was used to analyze fragments and a threshold of 300 relative fluorescent units (RFU) was used to call peaks from background noise. Decimals on size fragment calls were rounded.

### Long read amplicon sequencing of *msp2* and *in silico* fragment size analysis

Sequencing of *msp2* was performed by CCS as previously described [28]. Briefly, the entire open reading frame of *msp2* was amplified including start and stop codons, where each 15µl reaction contained 6 units Phusion Hot Start DNA Polymerase (Fischer Scientific), 200µM dNTP mix (Fischer Scientific), 0.5µM each forward and reverse primers (Fischer Scientific) for *msp2*. A nested 50µl reaction was then used to asymmetrically barcode *msp2* amplicons using barcoded forward and reverse primers. Nested reactions used 20 units of Phusion Hot Start II DNA Polymerase (Fischer Scientific), 0.5 μM each forward and reverse primers (Fischer Scientific), and 1μl template (product from the first PCR) per isolate. Pooled, cleaned amplicon pools were sequenced (CCS) in SMRTcell™ 1M by PacBio Sequel I with Sequel 3.0 polymerase. During the ongoing study, samples in the final 384-plex library were switched to PacBio Sequel II due to the updated technology. Read volume and noise of Sequel I and II were comparable. Reads from CCS were genotyped *in silico* using a pipeline in the European Galaxy server [28]. Briefly, the forward and reverse primers used for genotyping by the CE method were run as BLASTn queries against the CCS reads. A threshold for noise was determined using the calling rate for known reference *msp2* sequences. A false positivity rate (FPR) threshold of 0.01 was applied to both size variants and sequences.

### Accuracy in fragment sizing by CE and CCS based genotyping

To estimate the technical variability in CE genotyping, we assessed a library of mock infections created by mixing varying concentrations of known, reference strain *msp2* (**Figure S1**) obtained from PlasmoDB [28,68]: HB3 (LR131339 REGION: 250760.251530), CD01 (PfCD01_020011700), Dd2 (PfDd2_020009600), SN01 (PfS N01_020009800), KE01 (PfKE01_020009300), SD01 (PfSD01_020 012300), GN01 (PfGN01_020012100), 7G8 (Pf7G8_020011500), GB4 (PfGB4_020009800), and 3D7 (PF3D7_0206800). The difference between expected and observed sizes, as calculated *in silico* using oligos binding to the reference sequences, was determined (**Figure S1A**). The mock infections were submitted for CCS on Sequel I using the same protocol as described above. The difference between expected and observed *msp2* size variants by *in silico* genotyping was then calculated (**Figure S1B**). Reference sequences from PlasmoDB were aligned with called sequences using MAFFT FFT-NS-i (version 7.526) for all ten reference strains, to determine nucleotide errors (**Figure S2**).

### Data and statistical analysis

Size fragment data was analysed in R version 4.4.3. Statistical experts (NE, PV) performed binning and the statistical analysis of observed and expected CE size fragment prevalence (Cramér von Mises test). Diversity indices were evaluated using R packages “vegan”, “asbio”, and “adiv”. Rarefaction curves were created using the “vegan” R package. Mean prevalence of common variants in site populations was performed using two-tailed t test with the “rstatix” package. Comparison of mean MOI between transmission intensity groups was performed using Kruskal-Wallis test in R.

Nucleotide sequences were aligned using MAFFT in the European Galaxy server with Kimura 200 model and FFT-NS-i for slow, iterative alignment. Phylogenetic trees were created using the same parameters, including *P. billcollinsi* G01 *msp2* as the root. A maximum likelihood tree was created using IQ tree with 1000 ultrafast bootstrap replicates and 1000 replicates SH-like approximate likelihood ratio (SH-aLRT) single-branch tests. Maximum likelihood trees were visualized in iTOL. To determine protein sequence conservation, predicted protein sequences were aligned using MAFFT with BLOSUM 62 calculations and FFT-NS-2 for fast, progressive alignment. To evaluate *msp2* size fragment sequences, the *in silico* version of nested PCR primers (used prior to CE genotyping) were run against called sequence variants using BLASTn. The 5 prime- and 3 prime-most base pair location of primer binding sites were used to trim sequences using the Trim tool (Galaxy Version 0.0.2). A pairwise Hamming distance matrix was used to perform principal component analysis of block 3 sequences. Expected heterozygosity (H_e_) for full and size fragment *msp2* sequences was assessed by taking average H_e_ across all sequences belonging to the same size variant. Sequence alignments and primer binding locations were visualized in Geneious version 2024.0 created by Biomatters (available from https://www.geneious.com). Pairwise sequence identity matrices were calculated in Geneious after alignment using Clustal, and visualized in R using ggplot2.

### Term definitions

Size variants are defined as distinct *msp2* fragment sizes of either FC27 or IC type. Unique full length *msp2* sequences are the entire *msp2* open reading frame and are distinct from each other by at least one nucleotide. Unique size fragment sequences are defined as the region targeted by *in silico* PCR primers, including the primer binding regions (**Figure 1B**), and are distinct from others by at least one nucleotide. Unique amino acid sequences were predicted from unique full length nucleotide sequences and are defined by one amino acid difference from each other.

## Data availability

Fastq files have been uploaded onto the European Nucleotide Archive (ENA) at project number PRJEB91187. Fastq files for three of the travelers (Trav_Gambia_34 [ERS21325985], Trav_Ghana_31 [ERS21325986], Trav_Ghana_29 [ERS21325988]) and their respective controls are available at ENA project number PRJEB81353; for Trav_Uganda_4 [ERS25101145] and respective controls at project number PRJEB91187. Fastq files for mock infections are available at ENA project number PRJEB81353. The galaxy pipeline used for *in silico* genotyping is available at https://zenodo.org/record/8177047. Other data files can be made available upon request. Figures were created in BioRender and are available at https://BioRender.com/zau0t8e.

## Acknowledgement

We are grateful to all study participants and communities for their contributions. We thank all the research teams, investigators, and students involved in the included studies. We would like to thank Maria de Jesus Santos (CNE Health Ministry, São Tomé and Príncipe) for contributing with samples from São Tomé and Príncipe. We also greatly appreciate the Center for Molecular Medicine (CMM) for the use of their core facilities, and SciLifeLab (Uppsala) for the use of their PacBio Sequell I and II instruments.

## Author contributions

All authors contributed to the work presented in this paper, and accept responsibility for the included studies, data collection, and analysis. JZ, IB, gathered data and performed analysis including visualization; KMK gathered data, performed analysis including visualization, reviewed and edited the paper; DP performed analysis including visualization, reviewed and edited the paper; FF collected samples and provided technical support; CA, MKC, CF, MJS, KMK, MK, SMK, MK, AL, DL, DDM, JO, KS, TNS collected samples, gathered data and performed analysis for respective studies, reviewed and edited the paper; NE, PV provided statistical expertise and reviewed the paper; EA, HB, PB, MC, PDC, JPG, KM, BN, JN, KEMP, SP, UR, SBS, BT, MKT, LIOO, AF designed respective studies, reviewed and edited the paper; JZ, IB, DP, AF wrote the manuscript; DP, AF conceptualized and designed the study and provided funding.

## References

[1] World Health Organization. World malaria report 2024: addressing inequity in the global malaria response. Geneva: Licence: CC BY-NC-SA 3.0 IGO; 2024.

[2] Vreden SGS, Verhave JP, Oettinger T, Sauerwein RW, Meuwissen JHET. Phase I clinical trial of a recombinant malaria vaccine consisting of the circumsporozoite repeat region of Plasmodium falciparum coupled to hepatitis B surface antigen. American Journal of Tropical Medicine and Hygiene 1991;45:533–8. 10.4269/AJTMH.1991.45.533.

[3] Datoo MS, Natama MH, Somé A, Traoré O, Rouamba T, Bellamy D, et al. Efficacy of a low-dose candidate malaria vaccine, R21 in adjuvant Matrix-M, with seasonal administration to children in Burkina Faso: a randomised controlled trial. Lancet 2021;397:1809–18. 10.1016/S0140-6736(21)00943-0.

[4] Ladda R, Aikawa M, Sprinz H. Penetration of erythrocytes by merozoites of mammalian and avian malarial parasites. J Parasitol 1969;55:633–44. 10.2307/3277308.

[5] Sanders PR, Kats LM, Drew DR, O’Donnell RA, O’Neill M, Maier AG, et al. A Set of Glycosylphosphatidyl Inositol-Anchored Membrane Proteins of Plasmodium falciparum Is Refractory to Genetic Deletion. Infect Immun 2006;74:4330. 10.1128/IAI.00054-06.

[6] Richards JS, Beeson JG. The future for blood-stage vaccines against malaria. Immunol Cell Biol 2009;87:377–90. 10.1038/ICB.2009.27.

[7] Osier FH, Mackinnon MJ, Crosnier C, Fegan G, Kamuyu G, Wanaguru M, et al. Malaria: New antigens for a multicomponent blood-stage malaria vaccine. Sci Transl Med 2014;6. 10.1126/scitranslmed.3008705.

[8] Health Organization W. Informal consultation on methodology to distinguish reinfection from recrudescence in high malaria transmission areas Report of a virtual meeting, 17–18 May 2021 Global Malaria Programme n.d.

[9] Tadele G, Jaiteh FK, Oboh M, Oriero E, Dugassa S, Amambua-Ngwa A, et al. Low genetic diversity of Plasmodium falciparum merozoite surface protein 1 and 2 and multiplicity of infections in western Ethiopia following effective malaria interventions. Malar J 2022;21:383. 10.1186/s12936-022-04394-1.

[10] Prescott N, Stowers AW, Cheng Q, Bobogare A, Rzepczyk CM, Saul A. Plasmodium falciparum genetic diversity can be characterised using the polymorphic merozoite surface antigen 2 (MSA-2) gene as a single locus marker. Mol Biochem Parasitol 1994;63:203–12. 10.1016/0166-6851(94)90056-6.

[11] Messerli C, Hofmann NE, Beck HP, Felger I. Critical Evaluation of Molecular Monitoring in Malaria Drug Efficacy Trials and Pitfalls of Length-Polymorphic Markers. Antimicrob Agents Chemother 2016;61. 10.1128/AAC.01500-16.

[12] Hastings IM, Felger I. WHO antimalarial trial guidelines: good science, bad news? Trends Parasitol 2022;38:933–41. 10.1016/J.PT.2022.08.005.

[13] Snounou G, Beck HP. The use of PCR genotyping in the assessment of recrudescence or reinfection after antimalarial drug treatment. Parasitology Today 1998;14:462–7. 10.1016/S0169-4758(98)01340-4.

[14] Chen J-T, Li J, Zha G-C, Huang G, Huang Z-X, Xie D-D, et al. Genetic diversity and allele frequencies of Plasmodium falciparum msp1 and msp2 in parasite isolates from Bioko Island, Equatorial Guinea. Malar J 2018;17:458. 10.1186/s12936-018-2611-z.

[15] Oboh MA, Ndiaye T, Diongue K, Ndiaye YD, Sy M, Deme AB, et al. Allelic diversity of MSP1 and MSP2 repeat loci correlate with levels of malaria endemicity in Senegal and Nigerian populations. Malar J 2021;20. 10.1186/s12936-020-03563-4.

[16] Abukari Z, Okonu R, Nyarko SB, Lo AC, Dieng CC, Salifu SP, et al. The diversity, multiplicity of infection and population structure of P. Falciparum parasites circulating in asymptomatic carriers living in high and low malaria transmission settings of Ghana. Genes (Basel) 2019;10. 10.3390/genes10060434.

[17] Simpson S V., Nundu SS, Arima H, Kaneko O, Mita T, Culleton R, et al. The diversity of Plasmodium falciparum isolates from asymptomatic and symptomatic school-age children in Kinshasa Province, Democratic Republic of Congo. Malar J 2023;22:102. 10.1186/S12936-023-04528-Z.

[18] Amodu OK, Oyedeji SI, Ntoumi F, Orimadegun AE, Gbadegesin RA, Olumese PE, et al. Complexity of the msp2 locus and the severity of childhood malaria, in south-western Nigeria. Ann Trop Med Parasitol 2008;102:95–102. 10.1179/136485908X252340.

[19] Duah NO, Matrevi SA, Quashie NB, Abuaku B, Koram KA. Genetic diversity of Plasmodium falciparum isolates from uncomplicated malaria cases in Ghana over a decade malaria transmission seasons 2003. 10.1186/s13071-016-1692-1.

[20] Kimenyi KM, Wamae K, Ngoi JM, de Laurent ZR, Ndwiga L, Osoti V, et al. Maintenance of high temporal Plasmodium falciparum genetic diversity and complexity of infection in asymptomatic and symptomatic infections in Kilifi, Kenya from 2007 to 2018. Malar J 2022;21. 10.1186/s12936-022-04213-7.

[21] Mwingira F, Nkwengulila G, Schoepflin S, Sumari D, Beck HP, Snounou G, et al. Plasmodium falciparum msp1, msp2 and glurp allele frequency and diversity in sub-Saharan Africa. Malar J 2011;10. 10.1186/1475-2875-10-79.

[22] Ferreira MU, Hartl DL. Plasmodium falciparum: worldwide sequence diversity and evolution of the malaria vaccine candidate merozoite surface protein-2 (MSP-2). Exp Parasitol 2007;115:32–40. 10.1016/J.EXPPARA.2006.05.003.

[23] Smythe JA, Coppel RL, Day KP, Martin RK, Oduola AMJ, Kemp DJ, et al. Structural diversity in the Plasmodium falciparum merozoite surface antigen 2. Proceedings of the National Academy of Sciences 1991;88:1751–5. 10.1073/PNAS.88.5.1751.

[24] Snounou G. Genotyping of Plasmodium spp. Nested PCR. Methods Mol Med 2002;72:103–16. 10.1385/1-59259-271-6:103.

[25] Felger I, Snounou G, Hastings I, Moehrle JJ, Beck HP. PCR correction strategies for malaria drug trials: updates and clarifications. Lancet Infect Dis 2020;20:e20–5. 10.1016/S1473-3099(19)30426-8.

[26] Liljander A, Wiklund L, Falk N, Kweku M, Mrtensson A, Felger I, et al. Optimization and validation of multi-coloured capillary electrophoresis for genotyping of Plasmodium falciparum merozoite surface proteins (msp1 and 2). Malar J 2009;8:1–14. 10.1186/1475-2875-8-78/TABLES/4.

[27] Broumou I, Plaza DF, Färnert A. Genotyping of Plasmodium falciparum to Assess Clone Composition in Parasite Cultures. Methods in Molecular Biology 2022;2470:51–68. 10.1007/978-1-0716-2189-9_6/COVER.

[28] Plaza DF, Zerebinski J, Broumou I, Lautenbach MJ, Ngasala B, Sundling C, et al. A genomic platform for surveillance and antigen discovery in Plasmodium spp. using long-read amplicon sequencing. Cell Reports Methods 2023;3:100574. 10.1016/J.CRMETH.2023.100574.

[29] de Cesare M, Mwenda M, Jeffreys AE, Chirwa J, Drakeley C, Schneider K, et al. Flexible and cost-effective genomic surveillance of P. falciparum malaria with targeted nanopore sequencing. Nature Communications 2024 15:1 2024;15:1–16. 10.1038/s41467-024-45688-z.

[30] Färnert A, Yman V, Homann MV, Wandell G, Mhoja L, Johansson M, et al. Epidemiology of malaria in a village in the Rufiji River Delta, Tanzania: Declining transmission over 25 years revealed by different parasitological metrics. Malar J 2014;13:1–12. 10.1186/1475-2875-13-459/FIGURES/6.

[31] Andrade CM, Fleckenstein H, Thomson-Luque R, Doumbo S, Lima NF, Anderson C, et al. Increased circulation time of Plasmodium falciparum underlies persistent asymptomatic infection in the dry season. Nat Med 2020;26:1929–40. 10.1038/s41591-020-1084-0.

[32] Chen Y-A, Ng P-Y, Garcia-Ruiz D, Elliot A, Palmer B, Mendes Costa D’ R, et al. Genetic surveillance reveals low but sustained malaria transmission with clonal replacement in Sao Tome and Principe. Communications Medicine 2025 5:1 2025;5:1–12. 10.1038/s43856-025-00905-8.

[33] Chen YA, Shiu TJ, Tseng LF, Cheng CF, Shih WL, de Assunção Carvalho AV, et al. Dynamic changes in genetic diversity, drug resistance mutations, and treatment outcomes of falciparum malaria from the low-transmission to the pre-elimination phase on the islands of São Tomé and Príncipe. Malar J 2021;20:467. 10.1186/S12936-021-04007-3.

[34] Arez AP, Snounou G, Pinto J, Sousa CA, Modiano D, Ribeiro H, et al. A clonal Plasmodium falciparum population in an isolated outbreak of malaria in the Republic of Cabo Verde. Parasitology 1999;118:347–55. 10.1017/S0031182099003972.

[35] Girgis ST, Adika E, Nenyewodey FE, Senoo Jnr DK, Ngoi JM, Bandoh K, et al. Drug resistance and vaccine target surveillance of Plasmodium falciparum using nanopore sequencing in Ghana. Nature Microbiology 2023 8:12 2023;8:2365–77. 10.1038/s41564-023-01516-6.

[36] Oraby AK, Stojic A, Elawar F, Bilawchuk LM, McClelland RD, Erwin K, et al. A single amino acid mutation alters multiple neutralization epitopes in the respiratory syncytial virus fusion glycoprotein. Npj Viruses 2025;3:33. 10.1038/s44298-025-00119-8.

[37] Zhong D, Afrane Y, Githeko A, Yang Z, Cui L, Menge DM, et al. Plasmodium falciparum Genetic Diversity in Western Kenya Highlands. Am J Trop Med Hyg 2007;77:1043–50. 10.4269/AJTMH.2007.77.1043.

[38] Flück C, Schöpflin S, Smith T, Genton B, Alpers MP, Beck H-P, et al. Effect of the malaria vaccine Combination B on merozoite surface antigen 2 diversity 2006. 10.1016/j.meegid.2006.03.006.

[39] Genton B, Al-Yaman F, Anders R, Saul A, Brown G, Pye D, et al. Safety and immunogenicity of a three-component blood-stage malaria vaccine in adults living in an endemic area of Papua New Guinea. Vaccine 2000;18:2504–11. 10.1016/S0264-410X(00)00036-0.

[40] Genton B, Al-Yaman F, Betuela I, Anders RF, Saul A, Baea K, et al. Safety and immunogenicity of a three-component blood-stage malaria vaccine (MSP1, MSP2, RESA) against Plasmodium falciparum in Papua New Guinean children. Vaccine 2003;22:30–41. 10.1016/S0264-410X(03)00536-X.

[41] Genton B, Betuela I, Felger I, Al-Yaman F, Anders RF, Saul A, et al. A Recombinant Blood-Stage Malaria Vaccine Reduces Plasmodium falciparum Density and Exerts Selective Pressure on Parasite Populations in a Phase 1-2b Trial in Papua New Guinea. J Infect Dis 2002;185:820–7. 10.1086/339342.

[42] Patarroyo ME, Aza-Conde J, Moreno-Vranich A, Pabón L, Varela Y, Patarroyo MA. Far from the madding crowd: The molecular basis for immunological escape of plasmodium falciparum. Curr Issues Mol Biol 2017;22:65–78. 10.21775/cimb.022.065.

[43] Molina-Franky J, Patarroyo ME, Kalkum M, Patarroyo MA. The Cellular and Molecular Interaction Between Erythrocytes and Plasmodium falciparum Merozoites. Front Cell Infect Microbiol 2022;12. 10.3389/FCIMB.2022.816574.

[44] Ranford-Cartwright LC, Taylor RR, Asgari-Jirhandeh N, Smith DB, Roberts PE, Robinson VJ, et al. Differential antibody recognition of FC27-like Plasmodium falciparum merozoite surface protein MSP2 antigens which lack 12 amino acid repeats. Parasite Immunol 1996;18:411–20. 10.1046/J.1365-3024.1996.D01-137.X.

[45] Tonhosolo R, Wunderlich G, Ferreira MU. Differential Antibody Recognition of Four Allelic Variants of the Merozoite Surface Protein-2 (MSP-2) of Plasmodium falciparum. Journal of Eukaryotic Microbiology 2001;48:556–64. 10.1111/J.1550-7408.2001.TB00191.X.

[46] Tonon AP, Hoffmann EHE, Silveira LA Da, Ribeiro AG, Gonçalves CRDS, Ribolla PEM, et al. Plasmodium falciparum: sequence diversity and antibody recognition of the Merozoite surface protein-2 (MSP-2) in Brazilian Amazonia. Exp Parasitol 2004;108:114–25. 10.1016/J.EXPPARA.2004.08.001.

[47] Zerebinski J, Margerie L, Han NS, Moll M, Ritvos M, Jahnmatz P, et al. Naturally acquired IgG responses to Plasmodium falciparum do not target the conserved termini of the malaria vaccine candidate Merozoite Surface Protein 2. Front Immunol 2024;15:1501700. 10.3389/FIMMU.2024.1501700/BIBTEX.

[48] Das SC, Morales RAV, Seow J, Krishnarjuna B, Dissanayake R, Anders RF, et al. Lipid interactions modulate the structural and antigenic properties of the C-terminal domain of the malaria antigen merozoite surface protein 2. FEBS Journal 2017;284:2649–62. 10.1111/febs.14135.

[49] Zhang X, Perugini MA, Yao S, Adda CG, Murphy VJ, Low A, et al. Solution conformation, backbone dynamics and lipid interactions of the intrinsically unstructured malaria surface protein MSP2. J Mol Biol 2008;379:105–21. 10.1016/J.JMB.2008.03.039.

[50] Reddy SB, Anders RF, Beeson JG, Färnert A, Kironde F, Berenzon SK, et al. High affinity antibodies to plasmodium falciparum merozoite antigens are associated with protection from malaria. PLoS One 2012;7. 10.1371/journal.pone.0032242.

[51] Reddy SB, Anders RF, Cross N, Mueller I, Senn N, Stanisic DI, et al. Differences in affinity of monoclonal and naturally acquired polyclonal antibodies against Plasmodium falciparum merozoite antigens. BMC Microbiol 2015;15. 10.1186/s12866-015-0461-1.

[52] Lu C, Zheng X, Zhang W, Zhao H, MacRaild CA, Norton RS, et al. Interaction of merozoite surface protein 2 with lipid membranes. FEBS Lett 2019;593:288–95. 10.1002/1873-3468.13320.

[53] MacRaild CA, Pedersen ME, Anders RF, Norton RS. Lipid interactions of the malaria antigen merozoite surface protein 2. Biochim Biophys Acta Biomembr 2012;1818:2572–8. 10.1016/j.bbamem.2012.06.015.

[54] Roy SW, Ferreira MU, Hartl DL. Evolution of allelic dimorphism in malarial surface antigens. Heredity (Edinb) 2008;100:103–10. 10.1038/sj.hdy.6800887.

[55] Adda CG, Murphy VJ, Sunde M, Waddington LJ, Schloegel J, Talbo GH, et al. Plasmodium falciparum merozoite surface protein 2 is unstructured and forms amyloid-like fibrils. Mol Biochem Parasitol 2009;166:159–71. 10.1016/j.molbiopara.2009.03.012.

[56] Vacic V, Oldfield CJ, Mohan A, Radivojac P, Cortese MS, Uversky VN, et al. Characterization of molecular recognition features, MoRFs, and their binding partners. J Proteome Res 2007;6:2351–66. 10.1021/PR0701411/ASSET/IMAGES/LARGE/PR0701411F00006.JPEG.

[57] Fuxreiter M, Tompa P, Simon I. Local structural disorder imparts plasticity on linear motifs. Bioinformatics 2007;23:950–6. 10.1093/BIOINFORMATICS/BTM035.

[58] Abeln S, Frenkel D. Disordered Flanks Prevent Peptide Aggregation. PLoS Comput Biol 2008;4:e1000241. 10.1371/JOURNAL.PCBI.1000241.

[59] Zhang W, Zhang J, MacRaild CA, Norton RS, Anders RF, Zhang X. Modulation of the aggregation of an amyloidogenic sequence by flanking-disordered region in the intrinsically disordered antigen merozoite surface protein 2. European Biophysics Journal 2019;48:99–110. 10.1007/S00249-018-1337-8/FIGURES/8.

[60] Bhattacherjee A, Wallin S. Coupled Folding-Binding in a Hydrophobic/Polar Protein Model: Impact of Synergistic Folding and Disordered Flanks. Biophys J 2012;102:569–78. 10.1016/J.BPJ.2011.12.008.

[61] Krishnarjuna B, Sugiki T, Morales RAV, Seow J, Fujiwara T, Wilde KL, et al. Transient antibody-antigen interactions mediate the strain-specific recognition of a conserved malaria epitope. Communications Biology 2018 1:1 2018;1:1–10. 10.1038/s42003-018-0063-1.

[62] Zhang M, Wang C, Otto TD, Oberstaller J, Liao X, Adapa SR, et al. Uncovering the essential genes of the human malaria parasite Plasmodium falciparum by saturation mutagenesis. Science (1979) 2018;360. 10.1126/science.aap7847.

[63] Meyer CG, May J, Arez AP, Gil JP, Do Rosario V. Review: Genetic diversity of Plasmodium falciparum: asexual stages. Tropical Medicine & International Health 2002;7:395–408. 10.1046/J.1365-3156.2002.00875.X.

[64] Yman V, White MT, Asghar M, Sundling C, Sondén K, Draper SJ, et al. Antibody responses to merozoite antigens after natural Plasmodium falciparum infection: Kinetics and longevity in absence of re-exposure. BMC Med 2019;17:1–14. 10.1186/S12916-019-1255-3/FIGURES/5.

[65] Färnert A, Bronner U, Lebbad M, Björkman A. POLYCLONAL PLASMODIUM FALCIPARUM MALARIA IN TRAVELERS AND SELECTION OF ANTIFOLATE MUTATIONS AFTER PROGUANIL PROPHYLAXIS. 2002.

[66] Wångdahl A, Wyss K, Saduddin D, Bottai M, Ydring E, Vikerfors T, et al. Severity of Plasmodium falciparum and Non-falciparum Malaria in Travelers and Migrants: A Nationwide Observational Study over 2 Decades in Sweden. Journal of Infectious Diseases 2019;220:1335–45. 10.1093/infdis/jiz292.

[67] Yman V, Wandell G, Mutemi DD, Miglar A, Asghar M, Hammar U, et al. Persistent transmission of Plasmodium malariae and Plasmodium ovale species in an area of declining Plasmodium falciparum transmission in eastern Tanzania. PLoS Negl Trop Dis 2019;13. 10.1371/JOURNAL.PNTD.0007414.

[68] Bahl A, Brunk B, Coppel RL, Crabtree J, Diskin SJ, Fraunholz MJ, et al. PlasmoDB: The Plasmodium genome resource. An integrated database providing tools for accessing, analyzing and mapping expression and sequence data (both finished and unfinished). Nucleic Acids Res 2002;30:87–90. 10.1093/nar/30.1.87.

