## Supplementary Tables for "Uncovering the genetic diversity of the malaria parasite antigen MSP2 across Sub-Saharan Africa"

**Table S1. Origin of *Plasmodium falciparum* isolates from 19 study sites across sub-Saharan Africa.** Description of *P.* *falciparum* isolates from 20 cohorts assessed by *msp2* genotyping using nested PCR and fragment sizing by capillary electrophoresis (CE) (n=2761), with a subset selected for analysis by high fidelity long-read circular consensus sequencing (CCS) (n=1317) on the PacBio Sequell platform.

|  | **Country** | **Study site** | **Reference** | **Study**  **design** | **Collection**  **years** | **Transmission level year of sampling**  **(*Pf*PR_2-10_)^a^** | **Type of cases** | **Age**  **range,**  years | **Sample material** | **DNA extraction method** | **CE^b^**  **data** | **Selection**  **CE pos**  **1 per individual^c^** | **Selection CCS** | **CE**  **N=2761** | **CCS**  **N=1317** |
| --- | --- | --- | --- | --- | --- | --- | --- | --- | --- | --- | --- | --- | --- | --- | --- |
| 1 | Angola | Mabubas | Fançony 2025 (1) | Drug trial | 2020-2021 | Moderate | Sympt | 2-10 | DBS | QIAamp DNA mini | N | Baseline | Micr pos | 99 | 99 |
| 2 | Burkina Faso | Tanghin Wobdo | Cherif in prep | Cohort study | 2020 | Moderate | Asympt | 0.5–5 | DBS | QIAamp DNA mini*** | N | First micr pos | Micr pos  CE pos | 82 | 81 |
| 3 | Ethiopia | Hawassa | Chaka in prep | Cohort study | 2016-2017 | Low | Sympt | 7-30 | DBS | QIAamp DNA mini | N | Baseline | CE pos | 23 | 23 |
| 4 | Ghana | Hohoe | Liljander 2011 (2) | IPTc trial | 2005-2006 | Moderate | Asympt | 0.3-5 | DBS | QIAamp DNA mini*** | Y | First pos | Micr pos | 544 | 83 |
| 5 | Kenya | Kilifi | Sondén 2013 (3) | Birth cohort | 2002-2006 | Low-moderate | Asympt+ sympt | 0-2 | Blood pellets | Puregene kits Qiagen | Y | First pos | nd | 33 | nd |
| 6 | Kenya | MALPAC | Khaemba 2020 (4) | Cohort study | 2013-2014 | Low | Asympt | 18-50 | Blood | QIAamp DNA mini kit | Y | First pos | nd | 84 | nd |
| 7 | Kenya | MVT | Liljander 2011 (5) | Vaccine trial | 2005-2006 | High | Asympt | 1-6 | Blood | ABI 6100 or Puregene kits | Y | First pos | nd | 369 | nd |
| 8 | Kenya | Junju | Kimenyi 2022 (6) | Cohort study | 2007-2018 | High-  Moderate | Asympt | 0-15 | Blood | QIAamp DNA mini kit | Y | First pos  +  All pos^e^ | CE sizes^d^ | 239 | 140 |
| 9 | Kenya | Busia | Nderu 2019 (7) | Drug trial | 2016 | Low | Sympt | 0.5-11 | Blood | QIAamp DNA mini kit | N | Baseline | nd | 79 | nd |
| 10 | Kenya | Msambweni | Nderu 2019 (7) | Drug trial | 2013 | Moderate | Sympt | 0.5-10 | DBS | QIAamp DNA mini kit | N | Baseline | nd | 35 | nd |
| 11 | Kenya | Nyando | Nderu 2019 (7) | Drug trial | 2015 | Low | Sympt | 0.7-11 | DBS | QIAamp DNA mini kit | N | Baseline | nd | 35 | nd |
| 12 | Mali | Kalifabougou | Portugal 2017 (8) | Cohort study | 2012 | High | Asympt | 9-10 | DBS | Qiagen DNA microAmp | N | First pos^f^ | Micr pos  availability | 17 | 22 |
|  | **Country** | **Study site** | **Reference** | **Study**  **design** | **Collection**  **years** | **Transmission level year of sampling**  **(*Pf*PR_2-10_)^a^** | **Type of cases** | **Age**  **range**  years | **Sample material** | **DNA extraction method** | **CE^b^ data available** | **Selection**  **CE pos**  **1 per individual^c^** | **Selection CCS** | **CE**  **N=2761** | **CCS**  **N=1317** |
| 13 | Mali | Kambila | Sondén 2015 (9) | Cohort study | 2006-2007 | High | Asympt | 2-25 | DBS | QIAamp DNA mini kit | Y | First pos | PCR pos | 139 | 59 |
| 14 | Nigeria | Ibadan | Tijani 2017 (10,11) | Cohort study | 2009-2010 | High | Sympt+  asympt | 0.5-43 | Blood | QIAamp DNA mini | N | CE  micr pos | Micr pos | 38 | 40 |
| 15 | Rwanda | Nyagatare | Surowiec 2015 (12) | Hospital study | 2011-2013 | Low | Sympt | 0.7-6 | Blood | QIAamp DNA mini | N | Baseline  Micr pos | All | 28 | 28 |
| 16 | São Tomé and Príncipe | São Tomé | Te, submitted | Surveillance | 2021 | Low | Sympt | nd | DBS | QIAamp DNA mini | N | All | All | 102 | 102 |
| 17 | Sudan | Asar | Babiker 1998 (13) | Cohort study | 1993 | Low | Sympt + asympt | nd | DBS | Chelex | N | All | All | 9 | 9 |
| 18 | Tanzania | Nyamisati | Yman 2019 (14) | Population  surveys | 1999, 2016 | High-  Moderate | Asympt | 1-84 | Blood | ABI 6100 1999 QIAamp DNA mini 2016 | Y 1999  N 2016 | CE pos | Micr pos | 319:  270 (1999)  49 (2016) | 156  (102,  54) |
| 19 | Uganda | Kampala | Kiwuwa 2013 (15) | Hospital study | 2007-2008 | Moderate | Sympt | 0.5-6 | Blood | Invitrogen Easy-DNA | Y | All | Availability | 117 | 114 |
| 20 | 33 SSA  countries | Sweden | Yman 2019 (16)  Wångdahl 2019 (17)  Wångdahl 2023 (18) | Traveller with malaria and  migrants screened | 1995-2023 | Mixed | Sympt+ asympt | 3-70 | Blood | QIAamp DNA mini | N | Baseline  CE pos | Micr or PCR pos availability | 370 | 361 |

Abbreviations: Asympt: asymptomatic, DBS: dried blood spots, CE: capillary electrophoresis, CCS: circular consensus sequencing, micr: microscopy, nd:no data/not done, pos: positive; sympt: symptomatic; Y: yes, N: No

a. Malaria transmission level estimated as *P. falciparum* parasite rate in children aged 2-10 years (*Pf*PR_2-10_) at the year of sampling in the respective study sites retrieved from Malaria Atlas Project (malariaatlas.org) based on latitude and longitude of the site

b. *msp2* genotyping by CE was performed with the same protocol (19), samples from Kenya cohorts were performed at KWTRP, Kilifi, Kenya and the remaining at KI in Stockholm, Sweden. For sample sets with CE data not available (N), annealing temperature was modified from 58 to 61֯C in the nested PCR annealing step

c. Baseline refers to sample before treatment, first pos refers to first CE positive sample in sample series from participants samples over time

d. Samples from Kenya Junju which were CCS sequenced and genotyped *in silico* were a selection of samples with common size variants identified by CE.

e. All annual surveys from Junju cohort, Kenya were only included in analyses over time (Figure x) other spatial analyses included only one sample per individual.

f. Availability of DNA

Data from published studies was included based on PCR positivity or positivity by *msp2* genotyping by capillary electrophoresis (CE).

**Table S2.** Observed and expected cumulative prevalences for the ten most commonly detected FC27 and IC size variants, per study site, assuming expected random distribution.

|  | Study site (N samples) | N samples (CE) | N observed FC27 alleles | Proportion (%) of FC27 alleles  Observed / expected (p) | N observed IC alleles | Proportion (%) of IC alleles  Observed / expected (p) |
| --- | --- | --- | --- | --- | --- | --- |
|  | Angola Mabubas | 99 | 110 | 78.2/ 48.6 (<0.001) | 149 | 36.9 / 30.8 (0.321) |
|  | Burkina Faso Tanghin Wobdo | 82 | 48 | 72.9 / 69.5; (0.817) | 110 | 42.7 / 35.9; (0.342) |
|  | Ethiopia Hawassa | 23 | 28 | 100 / 100; (-) | 6 | 100 / 100; (-) |
|  | Ghana Hohoe | 544 | 483 | 95.2 / 43.4; (<0.001) | 806 | 32.8 / 17.6; (<0.001) |
| Kenya KBC, Kilifi | | 33 | 23 | 87 / 95.5; (0.604) | 26 | 57.7 / 80.4; (0.064) |
| Kenya MALPAC, Junju | | 84 | 60 | 95 / 89.2; (0.487) | 92 | 55.4 / 45.2; (0.241) |
|  | Kenya MVT | 369 | 396 | 91.7 / 45.5; (<0.001) | 463 | 41.7 / 20.7; (<0.001) |
|  | Kenya Junju 2007-2013 | 194 | 160 | 58.8 / 36.7; (<0.001) | 161 | 41 / 30.6; (0.062) |
|  | Kenya Junju 2013-2018 | 45 | 119 | 70.6 / 48; (<0.001) | 118 | 33.9 / 34.4; (1) |
|  | Kenya Busia | 79 | 63 | 81 / 69.4; (0.21) | 79 | 35.4 / 42.4; (0.513) |
|  | Kenya Msambweni | 35 | 25 | 88 / 94.8; (0.608) | 45 | 57.8 / 63.5; (0.669) |
|  | Kenya Nyando | 35 | 35 | 68.6 / 77.1; (0.588) | 49 | 38.8 / 54.4; (0.156) |
|  | Mali Kalifabougou | 17 | 16 | 87.5 / 99.4; (0.49) | 16 | 62.5 / 96; (0.085) |
|  | Mali Kambila | 139 | 80 | 88.8 / 70.9; (0.008) | 164 | 43.3 / 30.2; (0.024) |
|  | Nigeria Igbo-Ora | 38 | 25 | 96 / 98.8; (1) | 38 | 42.1 / 64.8; (0.064) |
|  | Rwanda Nyagatare | 28 | 31 | 80.6 / 86.9; (0.733) | 41 | 51.2 / 64.5; (0.375) |
|  | São Tomé and Principe | 102 | 82 | 80.5 / 62.2; (0.019) | 105 | 100 / 100; (-) |
|  | Sudan Asar | 9 | 4 | 100 / 100; (-) * | 8 | 100 / 100; (-) * |
|  | Tanzania Nyamisati 1999 | 280 | 215 | 82.3 / 44; (<0.001) | 337 | 34.7 / 21.7; (<0.001) |
|  | Tanzania Nyamisati 2016 | 49 | 48 | 79.2 / 74.2; (0.812) | 98 | 32.7 / 38.4; (0.456) |
|  | Uganda Kampala | 123 | 96 | 88.5 / 67.3; (0.001) | 99 | 39.4 / 39.9; (1) |
|  | Travelers | 370 | 389 | 63.2 / 22.7; (<0.001) | 507 | 26.6 / 16.3; (<0.001) |

* Site with small number of samples

P-values from Cramér von Mises test in parenthesis.

**Table S3. Successful *msp2* genotyping and sequencing in the respective study sites**. A subset of the 2761 samples genotyped for *msp2* by nested PCR and fragment sizing by capillary electrophoresis (CE), were analysed by long-read sequencing (CCS) and *in silico* genotyping.

| Study Site | Samples sent for sequencing | Samples with CCS reads above noise (%) | Final sensitivity (%) | % polyclonal CE / CCS (size) / CCS (seq) |
| --- | --- | --- | --- | --- |
| Angola Mabubas | 73/99 | 80 (77) | 74 | 72.7 / 54.8 / 57.5 |
| Burkina Faso Tanghin Wobdo | 34/81 | 38 (47) | 42 | 50.0 / 26.5 / 32.4 |
| Ethiopia Hawassa | 15/23 | 17 (74) | 65 | 43.5 / 6.3 / 18.8 |
| Ghana Hohoe | 32/83 | 42 (51) | 39 | 70.8 / 34.4 / 40.6 |
| Kenya Busia | ND | - | - | 57.0 /- |
| Kenya Junju | 60/140 | 66 (47) | 43 | 63.2 /- * |
| Kenya MALPAC |  | -- |  | 50.0 /- |
| Kenya Msambweni | ND | - | - | 65.7 /- |
| Kenya MVT (Junju) | ND | - | - | 82.4 /- |
| Kenya KBC (Kilifi) | 0/0 | - | - | 27.3 /- |
| Kenya Nyando | 0/0 | - | - | 82.9 /- |
| Mali Kambila | 10/59 | 13 (22) | 17 | 41.0 / 20 / 20 |
| Mali Kalifabougou | 6/22 | 6 (27) | 27 | 70.6 / 50 / 50 |
| Nigeria Igbo-Ora | 8/40 | 10 (25) | 20 | 57.9 / 12.5 / 12.5 |
| Rwanda Nyagatare | 7/28 | 9 (32) | 25 | 85.7 / 42.9 / 42.9 |
| São Tomé and Principe | 78/102 | 80 (78) | 76 | 71.6 / 9.0 / 9.0 |
| Sudan Asar | 2/9 | 2 (22) | 22 | 44.4 / 0 / 0 ** |
| Tanzania Nyamisati | 109/156 [84/102; 25/54] | 119 (92) [90 (90)0; 29 (54)] | 70 [82, 46] | 61.1 [49.2, 11.9] / 45.0 [50,28] / 55.0 [60.7,36] |
| Uganda Kampala | 83/114 | 97 (85) | 73 | 41.0 / 61.4 / 68.7 |
| Travelers | 204/361 | 247 (68) | 57 | 73.5 / 29.4 / 34.3 |
|  | **721/1317** | **822 (62)** | **55** | **65.8/ 35.8 / 41.1** |

* Kenya Junju samples which were sequenced and genotyped *in silico* were a selection of samples with common size variants. Results from CCS are omitted because they do not accurately represent the study site.

** Only 2 samples of 9 from Sudan were successfully sequenced and genotyped *in silico*.

Brackets indicate values for the two survey years in Tanzania Nyamisati [1999, 2016].

**Table S4. Expected heterozygosity for *msp2* sequences belonging to the top 10 most prevalent size variants identified by CCS.** Average number of single nucleotide polymorphisms (SNPs) for nucleotide sequences corresponding to prevalent size variants (CCS) was used to calculate expected Heterozygosity (He). N, total detected sequences; Na, number of unique sequences.

| **Family** | **Size**  **variant** | **N (full)** | **Na (full)** | **SNPs (full)** | ***He* (full)** | **SNPs (B3)** | ***He* (size fragment)** |
| --- | --- | --- | --- | --- | --- | --- | --- |
| **FC27** | 328 | 124 | 42 | 39 | 0.18 | 28 | 0.23 |
|  | 364 | 119 | 26 | 42 | 0.22 | 30 | 0.29 |
|  | 292 | 107 | 29 | 23 | 0.25 | 23 | 0.25 |
|  | 376 | 62 | 5 | 19 | 0.25 | 10 | 0.39 |
|  | 256 | 51 | 21 | 21 | 0.21 | 16 | 0.23 |
|  | 220 | 48 | 22 | 27 | 0.21 | 21 | 0.21 |
|  | 436 | 16 | 6 | 29 | 0.34 | 28 | 0.33 |
|  | 472 | 15 | 2 | 7 | 0.22 | 6 | 0.23 |
|  | 301 | 13 | 2 | 95 | 0.15 | 89 | 0.14 |
|  | 412 | 12 | 5 | 17 | 0.29 | 13 | 0.32 |
| **IC** | 434 | 106 | 10 | 166 | 0.05 | 164 | 0.05 |
|  | 479 | 45 | 16 | 573 | 0.1 | 278 | 0.13 |
|  | 473 | 27 | 5 | 509 | 0.15 | 149 | 0.12 |
|  | 485 | 25 | 12 | 597 | 0.21 | 321 | 0.2 |
|  | 545 | 24 | 9 | 284 | 0.26 | 182 | 0.26 |
|  | 593 | 22 | 11 | 132 | 0.3 | 124 | 0.31 |
|  | 575 | 22 | 7 | 141 | 0.3 | 135 | 0.3 |
|  | 539 | 19 | 13 | 231 | 0.27 | 230 | 0.27 |
|  | 509 | 18 | 11 | 177 | 0.36 | 172 | 0.36 |
|  | 461 | 18 | 12 | 512 | 0.16 | 154 | 0.23 |

References

1.	Fançony C, Fortes-Gabriel E, Zage F, Alexiou E, Broumou I, Pernaute-Lau L, et al. Artemether-Lumefantrine treatment selects Plasmodium falciparum multidrug resistance 1 (pfmdr1) increased copy number among African malaria infections. J Infect Dis [Internet]. 2025 Mar 26 [cited 2025 Jun 5]; Available from: https://pubmed.ncbi.nlm.nih.gov/40138574/

2.	Liljander A, Chandramohan D, Kweku M, Olsson D, Montgomery SM, Greenwood B, et al. Influences of intermittent preventive treatment and persistent multiclonal Plasmodium falciparum infections on clinical malaria risk. PLoS One [Internet]. 2010 [cited 2025 Jun 5];5(10). Available from: https://pubmed.ncbi.nlm.nih.gov/21048970/

3.	Lundblom K, Murungi L, Nyaga V, Olsson D, Rono J, Osier F, et al. Plasmodium falciparum Infection Patterns Since Birth and Risk of Severe Malaria: A Nested Case-Control Study in Children on the Coast of Kenya. PLoS One. 2013 Feb 13;8(2).

4.	Khaemba EN, Ogwang C, Kinyanjui S, Muindi JM, Koske JK, Kimani D, et al. Comparing drug regimens for clearance of malaria parasites in asymptomatic adults using PCR in Kilifi County, Kenya: an open-label randomised controlled clinical trial (MalPaC). Wellcome Open Res. 2020 Feb 20;5:36.

5.	Liljander A, Bejon P, Mwacharo J, Kai O, Ogada E, Peshu N, et al. Clearance of asymptomatic P. falciparum Infections Interacts with the number of clones to predict the risk of subsequent malaria in Kenyan children. PLoS One [Internet]. 2011 Feb 24 [cited 2025 Jun 5];6(2):e16940–e16940. Available from: https://europepmc.org/articles/PMC3044709

6.	Kimenyi KM, Wamae K, Ngoi JM, de Laurent ZR, Ndwiga L, Osoti V, et al. Maintenance of high temporal Plasmodium falciparum genetic diversity and complexity of infection in asymptomatic and symptomatic infections in Kilifi, Kenya from 2007 to 2018. Malar J. 2022 Dec 1;21(1).

7.	Nderu D, Kimani F, Karanja E, Thiong’o K, Akinyi M, Too E, et al. Genetic diversity and population structure of Plasmodium falciparum in Kenyan–Ugandan border areas. Tropical Medicine and International Health [Internet]. 2019 May 1 [cited 2025 Jun 5];24(5):647–56. Available from: https://pubmed.ncbi.nlm.nih.gov/30816614/

8.	Portugal S, Tran TM, Ongoiba A, Bathily A, Li S, Doumbo S, et al. Treatment of Chronic Asymptomatic Plasmodium falciparum Infection Does Not Increase the Risk of Clinical Malaria Upon Reinfection. Clinical Infectious Diseases [Internet]. 2017 Mar 1 [cited 2022 May 11];64(5):645–53. Available from: https://academic.oup.com/cid/article/64/5/645/2739519

9.	Sonden K, Doumbo S, Hammar U, Vafa Homann M, Ongoiba A, Traord B, et al. Asymptomatic Multiclonal Plasmodium falciparum Infections Carried Through the Dry Season Predict Protection Against Subsequent Clinical Malaria. Journal of Infectious Diseases. 2015;212(4):608–16.

10.	Fasanya A, Mohammed N, Saleh BH, Tijani MK, Teleka A, Quintana M del P, et al. Anti-phosphatidylserine antibody levels are low in multigravid pregnant women in a malaria-endemic area in Nigeria, and do not correlate with anti-VAR2CSA antibodies. Front Cell Infect Microbiol. 2023;13.

11.	Oyewole TA, Mohammed NO, Osarenren BO, Tijani MK, Persson KEM, Falade MO. Plasmodium falciparum transmission based on merozoite surface protein 1 (msp1) and 2 (msp2) gene diversity and antibody responses in Ibadan, Nigeria. Parasite Epidemiol Control. 2024 Aug 1;26.

12.	Surowiec I, Orikiiriza J, Karlsson E, Nelson M, Bonde M, Kyamanwa P, et al. Metabolic signature profiling as a diagnostic and prognostic tool in pediatric Plasmodium falciparum malaria. Open Forum Infect Dis [Internet]. 2015 Apr 1 [cited 2025 Jun 5];2(2). Available from: https://pubmed.ncbi.nlm.nih.gov/26110164/

13.	Babiker HA, Abdel-Muhsin AMA, Ranford-Cartwright LC, Satti G, Walliker D. Characteristics of Plasmodium falciparum parasites that survive the lengthy dry season in eastern Sudan where malaria transmission is markedly seasonal. American Journal of Tropical Medicine and Hygiene [Internet]. 1998 [cited 2025 Jun 5];59(4):582–90. Available from: https://pubmed.ncbi.nlm.nih.gov/9790434/

14.	Yman V, Wandell G, Mutemi DD, Miglar A, Asghar M, Hammar U, et al. Persistent transmission of Plasmodium malariae and Plasmodium ovale species in an area of declining Plasmodium falciparum transmission in eastern Tanzania. PLoS Negl Trop Dis [Internet]. 2019 May 1 [cited 2022 Jun 7];13(5). Available from: /pmc/articles/PMC6555537/

15.	Kiwuwa MS, Ribacke U, Moll K, Byarugaba J, Lundblom K, Färnert A, et al. Genetic diversity of Plasmodium falciparum infections in mild and severe malaria of children from Kampala, Uganda. Parasitol Res [Internet]. 2013 Apr 14 [cited 2022 Jun 7];112(4):1691–700. Available from: https://link.springer.com/article/10.1007/s00436-013-3325-3

16.	Yman V, White MT, Asghar M, Sundling C, Sondén K, Draper SJ, et al. Antibody responses to merozoite antigens after natural Plasmodium falciparum infection: Kinetics and longevity in absence of re-exposure. BMC Med [Internet]. 2019 Jan 30 [cited 2023 Feb 28];17(1):1–14. Available from: https://bmcmedicine.biomedcentral.com/articles/10.1186/s12916-019-1255-3

17.	Wångdahl A, Wyss K, Saduddin D, Bottai M, Ydring E, Vikerfors T, et al. Severity of Plasmodium falciparum and Non-falciparum Malaria in Travelers and Migrants: A Nationwide Observational Study over 2 Decades in Sweden. Journal of Infectious Diseases [Internet]. 2019 Sep 13 [cited 2021 May 18];220(8):1335–45. Available from: /pmc/articles/PMC6743839/

18.	Wångdahl A, Bogale RT, Eliasson I, Broumou I, Faroogh F, Lind F, et al. Malaria parasite prevalence in Sub-Saharan African migrants screened in Sweden: a cross-sectional study. The Lancet Regional Health - Europe [Internet]. 2023 Apr 1 [cited 2025 Jun 5];27. Available from: https://pubmed.ncbi.nlm.nih.gov/37069854/

19.	Liljander A, Wiklund L, Falk N, Kweku M, Mrtensson A, Felger I, et al. Optimization and validation of multi-coloured capillary electrophoresis for genotyping of Plasmodium falciparum merozoite surface proteins (msp1 and 2). Malar J [Internet]. 2009 Apr 23 [cited 2022 Oct 14];8(1):1–14. Available from: https://malariajournal.biomedcentral.com/articles/10.1186/1475-2875-8-78
