## Supplementary Figures for "Uncovering the genetic diversity of the malaria parasite antigen MSP2 across Sub-Saharan Africa"

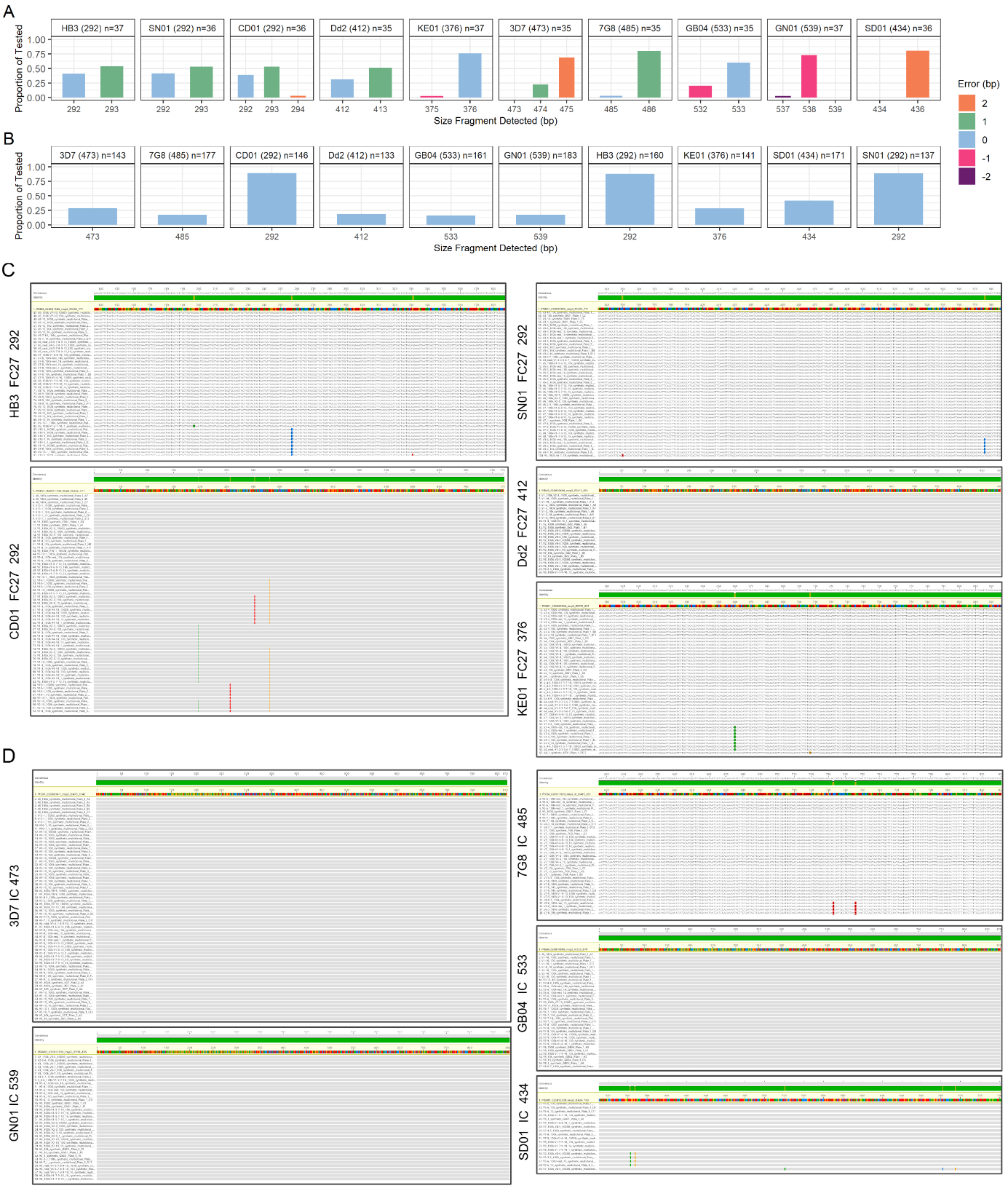

**Figure S1: Genotyping of mock infections by CE and CCS.** Mock infections were created by mixing varying concentrations of 1-10 known reference strain *msp2* sequences. Expected fragment sizes for each of the ten *msp2* variants and the proportion of tested mock infections in which the expected sizes were detected are shown for CE (**A**) and CCS (**B**) based genotyping. Size variants were not always detected, which is why not all facets make up 100% of tested mock infections with either method. Full length nucleotide sequences from CCS-based genotyping were aligned with the expected sequences, for FC27 type reference strain *msp2* (**C**) and IC type reference strain *msp2* (**D**). Sequences were aligned using MAFFT in Galaxy with FFT-NS-i for slow iterative refinement and visualized in Geneious. Divergence from expected reference strain sequences (highlighted) are shown with Clustal coloring.

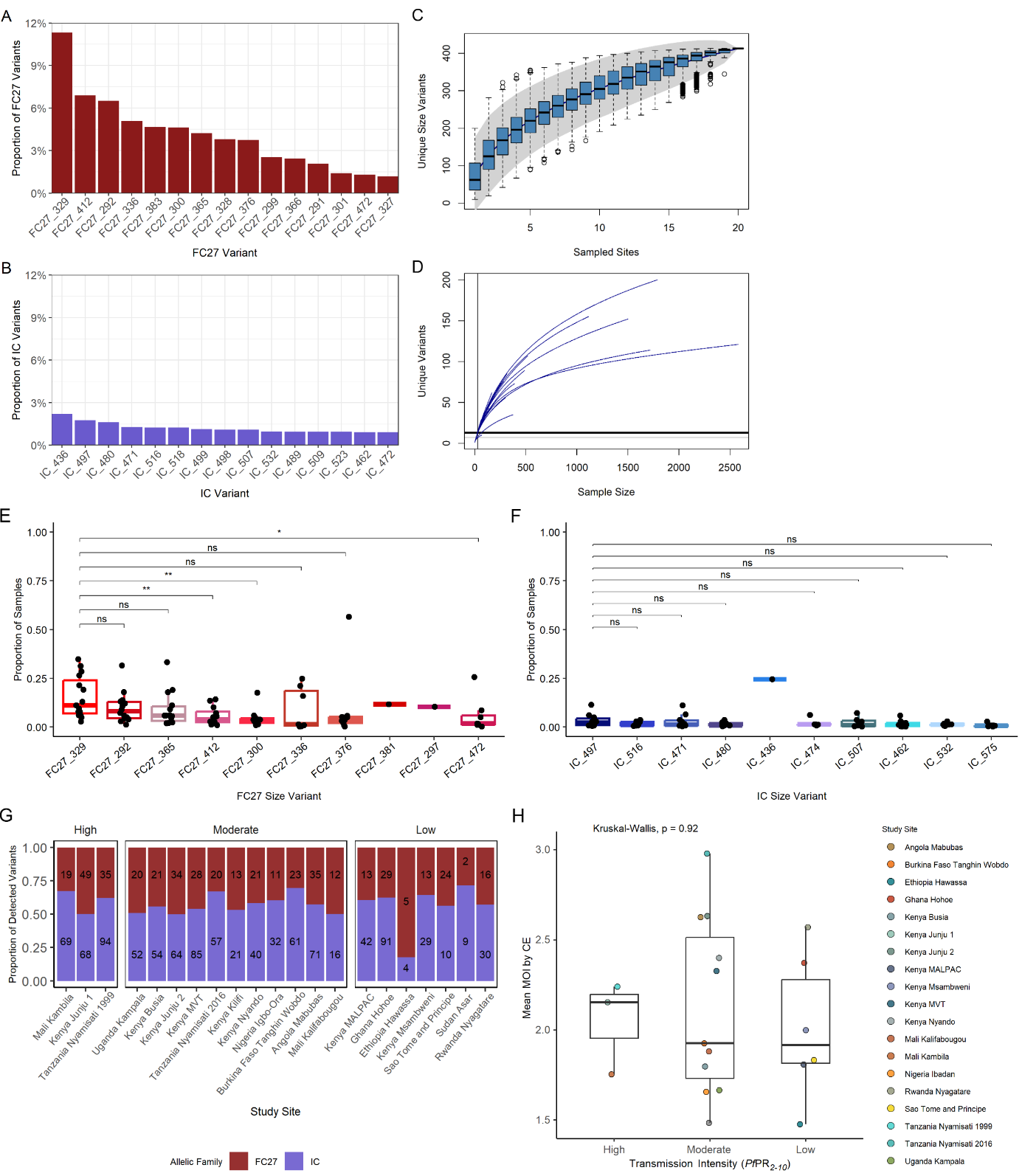

**Figure S2. Raw data from CE based genotyping shows patterns of prevalent *msp2* size variants.** Proportion of the 15 most common FC27 variants (**A**) and IC variants (**B**), prior to binning. Accumulation curve, modelled on binned data, for detected unique *msp2* size variants in relation to sites sampled (**C**). Model used 10000 permutations, and Lomolino method for extrapolating total number of species. Rarefaction curve for number of unique *msp2* variants with increasing sample size (**D**). Median proportion of samples carrying the 10 most prevalent FC27 size variants (**E**) and IC size variants (**F**) shown by boxplot. Proportion of samples between size variants compared using t-test; ns, not significant (p>0.05); * p<0.05, ** p<0.01.Proportion of detected *msp2* size variants belonging to the FC27 or IC family, at each site and grouped by transmission intensity (**G**). Text over columns identifies number of unique size variants per allelic family at the site. The median number of unique *msp2* alleles (MOI) per infection, for sites with low (1-10% *Pf*PR_2-10_), moderate (10-35% *Pf*PR_2-10_), or high transmission (>35% *Pf*PR_2-10_) reported by Malaria Atlas Project (**H**). Colored dots represent mean MOI at each site, mean MOI between transmission groups was compared using Kruskal-Wallis test.

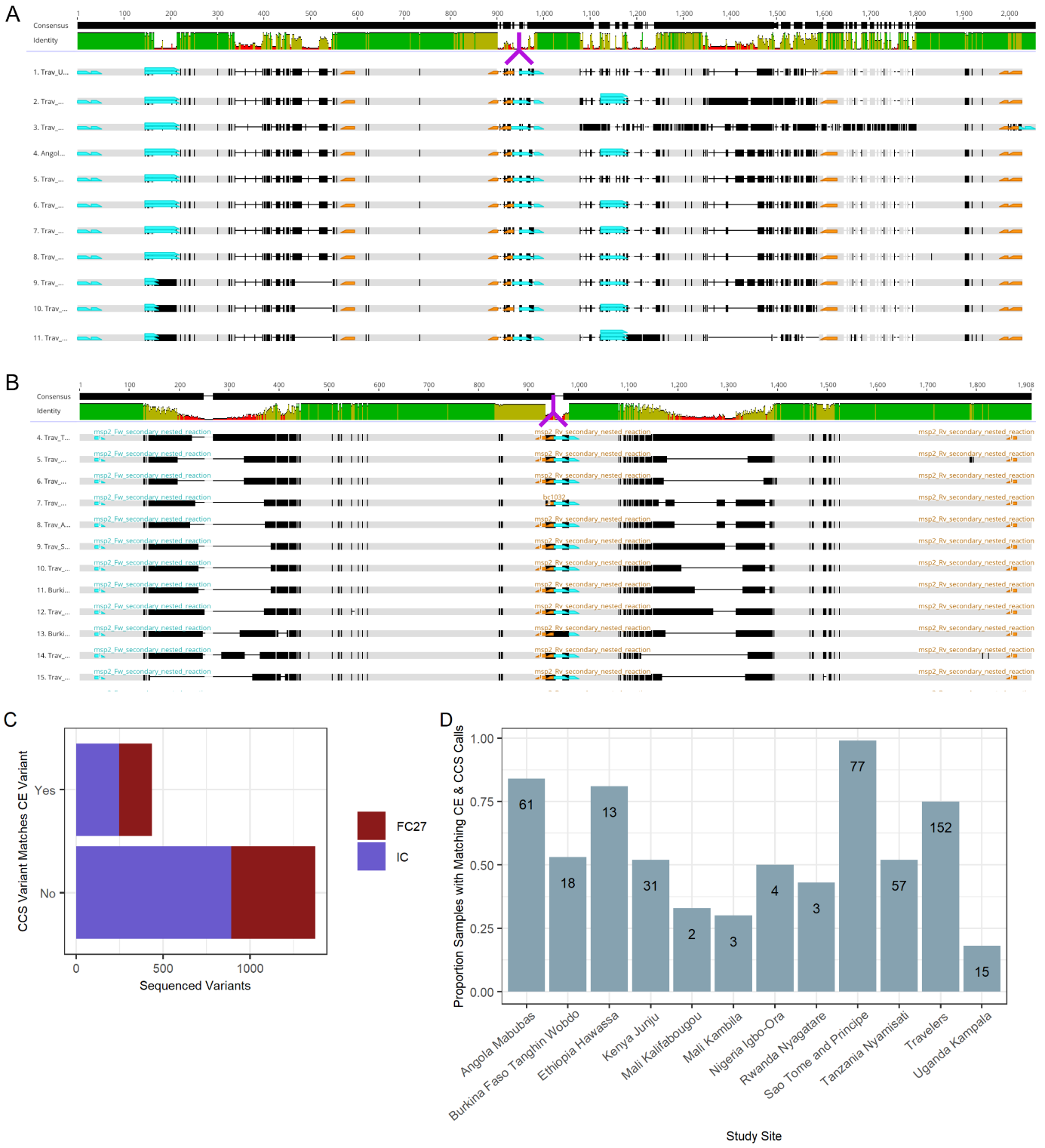

**Figure S3**. **Reads from long-read sequencing matching CE size variant** **calls.** Nucleotide sequences from the *in silico* genotyping pipeline were aligned using MAFFT and mapped in Geneious with primary and secondary nested PCR primers, barcode sequences from the secondary PCR reaction, and fragment size determining oligos. Forward primers and barcodes are indicated in teal and reverse primers and barcodes in orange. Sequences in (**A**) are a representative selection of FC27 *msp2* sequences determined to be artifacts, sequences in (**B**) are a representative selection of IC *msp2* sequences determined to be artifacts. Artifacts were determined by having two tandem *msp2* size fragments in addition to two sets of secondary reaction PCR primers with barcode sequences (shown by purple arrows) between the first reverse primer and the second forward primer. Sequences identified as artifacts were removed from further analysis. Sequences were determined to match CE size variants by having an *in silico* determined size variant within 3 base pairs length of one or more CE size variant call(s) from the same sample. Barplot shows the number of *msp2* sequences matching or not matching the CE call from the same sample (C). Proportion of samples from each study site with a matching call for CE and CCS (D), number of variants matching denoted in their respective bars.

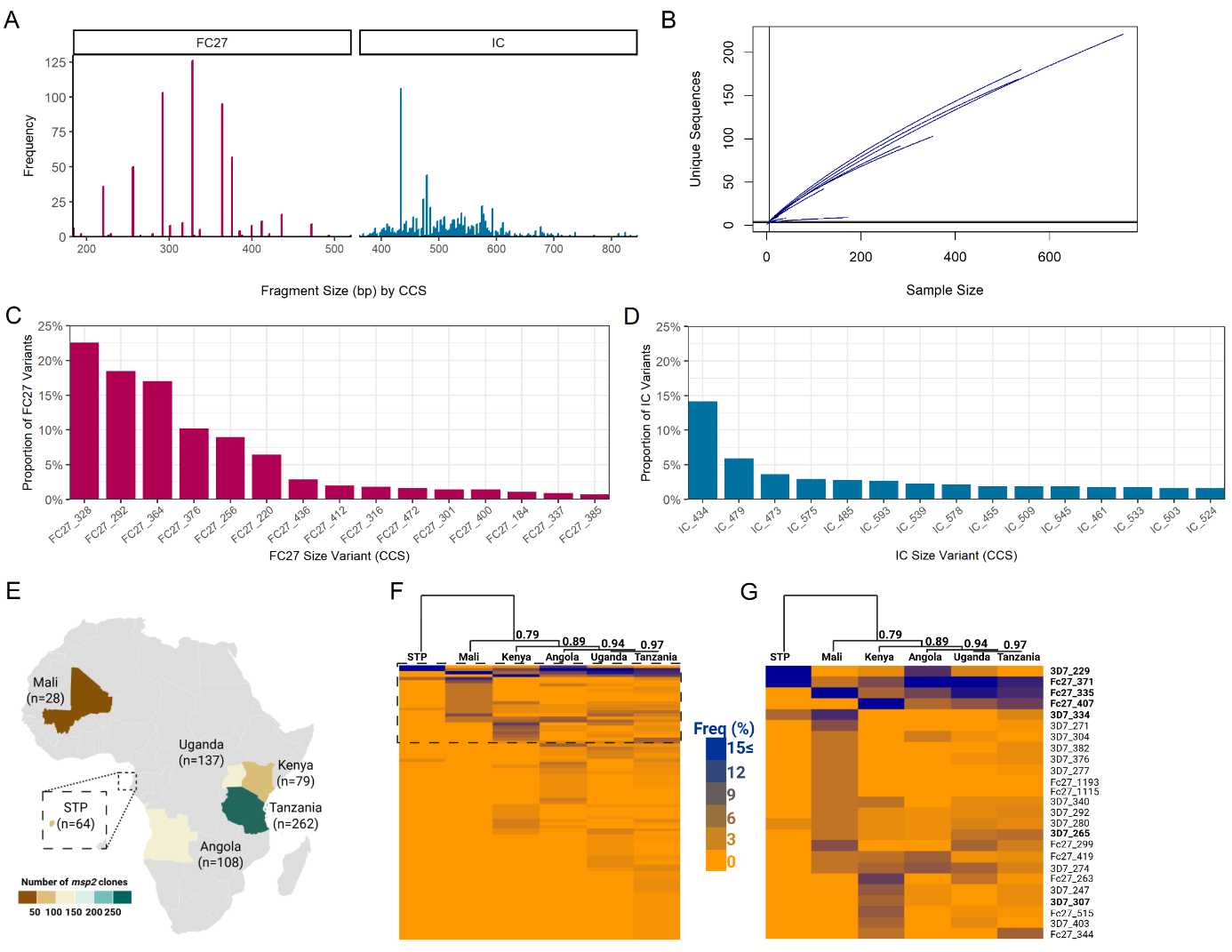

**Figure S4. CCS-based *in silico* genotyping identifies prevalent *msp2* variants.** Frequency of size variants identified using BLASTn against CCS reads (**A**). Rarefaction curve for unique full length *msp2* sequences by sample size (**B**). Proportion of the 15 most prevalent FC27 variants, out of all detected FC27 type *msp2* (**C**), and the 15 most prevalent IC variants, out of all detected IC type *msp2* (**D**). Size variants were determined *in silico* with a different set of primers used by Mwingira *et al* (1) in a subset of samples (n=678) sequenced for this study, including 153 Tanzania Nyamisati samples from additional surveys (**E**). Number of samples and country of isolate indicated by color bar. Frequency of Mwingira size variants in the sequences obtained for the 7 sites (**F**). The 91 Mwingira size variants identified in the sequencing dataset were hierarchically clustered (complete) by Euclidian distance metric. Countries were clustered using Pearson correlation (centered) as the similarity metric. A selection of the 25 Mwingira variants with the highest frequency in the sequencing dataset (dashed rectangle in **E**) is shown in (**G**). Variants shown by Mwingira *et al* to circulate with high frequencies (≥4%) in Burkina Faso, Sao Tome, Malawi, Tanzania and/or Uganda (1) indicated in bold.

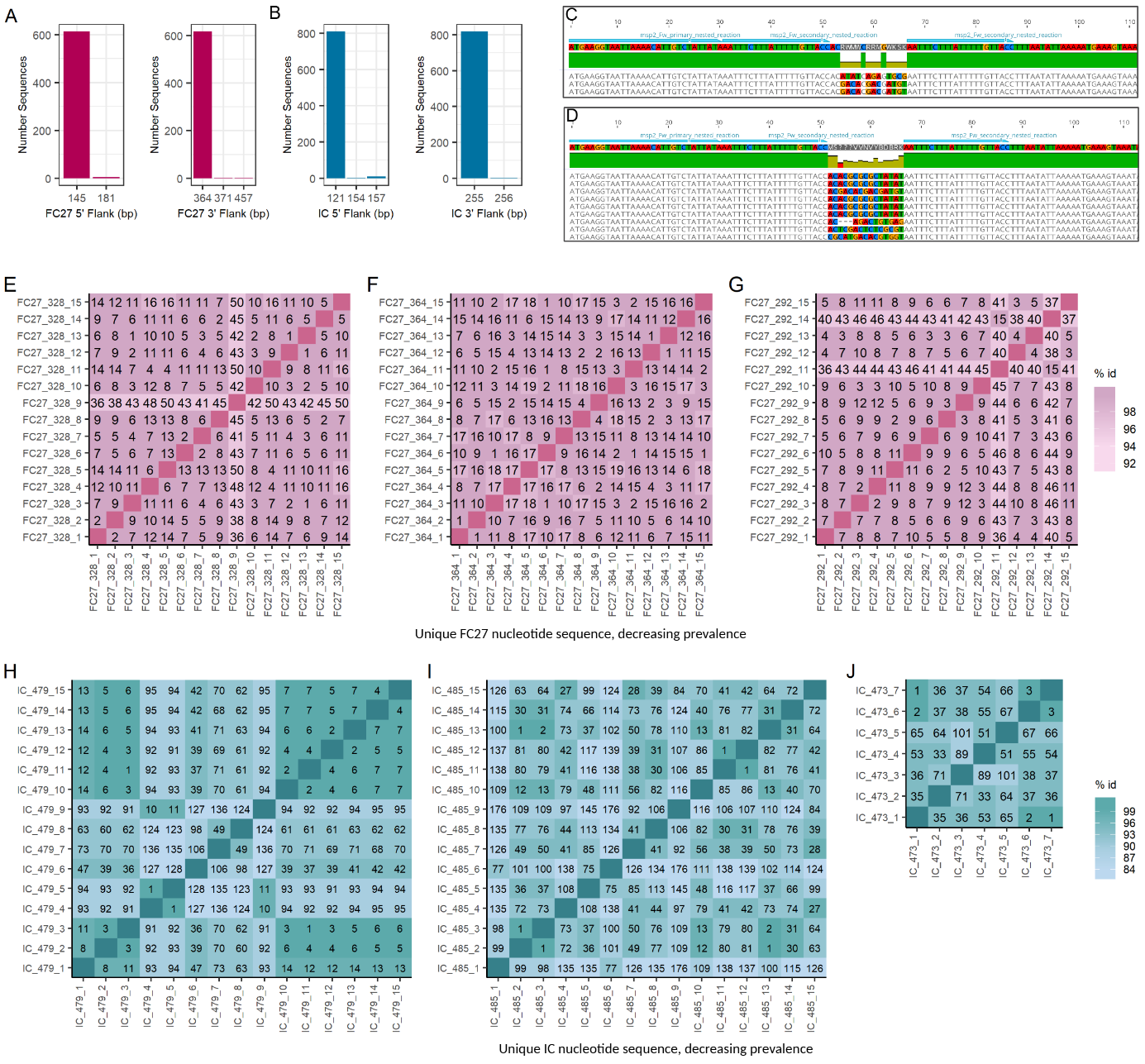

**Figure S5. Diversity in different regions of *msp2* nucleotide and predicted protein sequences.** The base pair length of the regions flanking the size fragment region was calculated for all msp2: upstream of the FC27 forward primer and downstream of the FC27 reverse primer (**A**), upstream of the IC forward primer and downstream of the IC reverse primer (**B**). Several sequence variants were identified as having insertions in block 1 for FC27 (**C**) and IC (**D**). Forward (turquoise) and reverse (orange) primers indicated. Bases not in agreement with consensus (95% threshold) highlighted in Clustal coloring. Pairwise similarity matrices for the top 15 most common, unique nucleotide sequences for FC27_328 (**E**), FC27_364 (**F**), FC27_292 (**G**), IC_479 (**H**), IC_485 (**I**), and all unique nucleotide sequences for IC_473 (**J**). Percent identity shown by color legend, base pair difference is overlaid in text. Sequences are named by decreasing prevalence, eg “FC27_328_1” the most common, “FC27_328_2” the second most common, and so on.

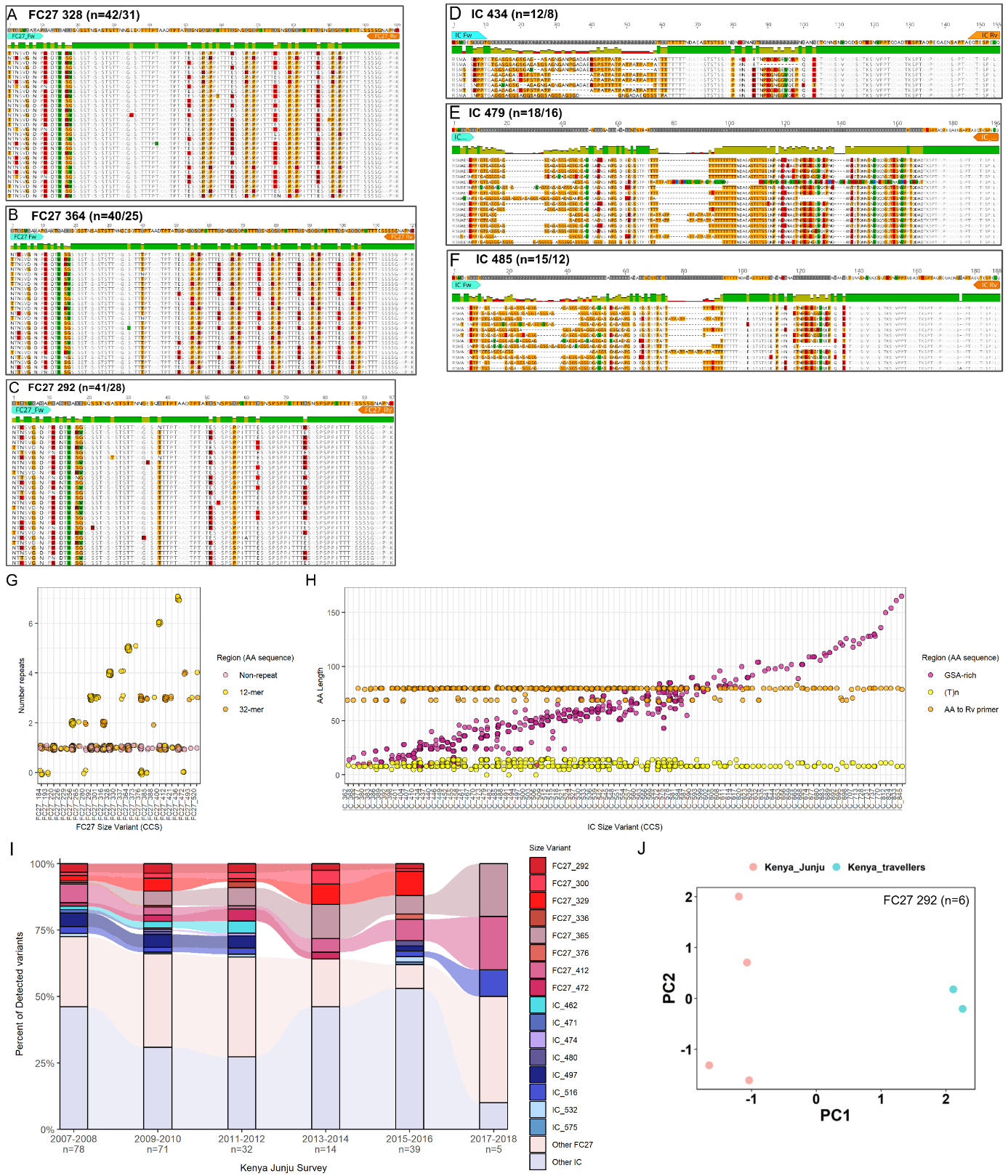

**Figure S6. Diversity in repeat regions for FC27 and IC amino acid sequences.** Unique, predicted amino acid sequences for each size variant detected by CCS and *in silico* genotyping were aligned using Clustal in Geneious. The size fragment determining region, identified using the secondary PCR forward (turquoise) and reverse (orange) oligos, is shown for FC27_328 (**A**), FC27_364 (**B**), FC27_292 (**C**), IC_434 (**D**), IC_479 (**E**), and IC_485 (**F**). Disagreement from consensus shown in Clustal coloring, number unique amino acid sequences and unique size fragment regions indicated in each panel. The number of FC27 family repeats, found within the block 3 region, for each size variant detected by CCS and *in silico* genotyping (**G**). The amino acid length of the GSA-rich region and tandem “T” repeats in the IC block 3 region, and the remaining length from the block 3 3’ end to the IC reverse primer (**H**). Distribution of CE-determined *msp2* size variants in samples from Kenya Junju without repeated sampling, with survey years grouped by two (**I**). Size fragment sequences belonging to FC27 292 (n=6) were plotted by PCA for Kenya Junju (coral) and Travelers to Kenya (teal) (**J**).

**Supplementary methods**

Genotyping of *msp2* using fluorescent family-specific primers, followed by fragment size analysis by CE, identifies length in base pairs. Lengths are identified to 2 decimal places and can vary between runs of known lengths, for example, the HB3 *msp2* size fragment (FC27 292) may have a peak anywhere between 291 and 293 (2). Genotyping often bins fragment sizes within 3 base pairs of each other (3–6), for *msp2* and other coding genes (7), and other loci with short tandem repeats (8).

FC27 *msp2* are known to increase in length due to tandem repeats of 12 amino acids, as well as repeats of 32 amino acids. IC *mps2,* on the other hand, increase in length due to additional GSA-rich repeats and T_n_ at the end of block 3, along with some variation in the non-repeat sequence in block 3 (9). The minimum length difference between IC fragments would be 3 nucleotides. By genotyping mock infections of known reference strain *msp2* sequences, we identified called fragment size with 1 bp less than or greater than expected length (**Figure S1A**)*.* The maximum difference was +2 bp (3D7 and SD01), but we felt that bins of greater than 3 bp would not be biologically relevant when applied to all size variants. The mean number of *msp2* genotypes per sample after binning (2.21) was not significantly different from the mean number detected before binning (2.26) by two-tailed t test (t = 1.2567, df = 5519.9, p value = 0.2089, 0.95 CI -0.025824--0.11497), which indicates binning did not significantly impact MOI measures in our data.

Size variants were determined from size fragment oligos using a published, accessible pipeline in Galaxy (10–12), with a false positivity rate of 0.01 for both size variant and sequence calling. CCS-determined size variants were not binned since results from mock infections did not identify any variation in variant length (**Figure S1B**). We did find sequences with incorrect bases (compared to expected) for several of the reference strain sequences (**Figure S1C,D**).

Transmission estimates were based on estimated parasite rates in children 2-10 years old (*Pf*PR_2-10_)from the Malaria Atlas Project (13,14). For study cohorts, which have known geographical locations, the site was used for the estimate. For travelers the exact location was not known, therefore these samples were not used for transmission analyses. Visualization of figures used R packages ggplot2(15), ggpubr (16), patchwork (17), scales (18), and ggdist (19,20). Diversity index analyses were performed using R version 4.4.3 with packages vegan (21), asbio (22), ggalluvial (23), and adiv (24). Rarefaction curve was created using vegan (21). Nucleotide sequences were aligned using MAFFT (FFT-NS-i) in Galaxy (25–27) and visualized in Geneious version 2024.0 created by Biomatters (available from [https://www.geneious.com](https://www.geneious.com/)). Phylogenetic trees were constructed using MAFFT (FFT-NS-i) in Galaxy of full length *msp2* and *msp2* from *P. billcollinsi* G01 available in PlasmoDB (28–31), followed by IQtree (32–35) with 1000 ultrafast bootstrap replicates (32,36) and SH-aLRT single branch tests (37) and visualized in Interactive Tree of Life (iTOL) (38,39) with root at *P. billcollinsi* G01 *msp2.*

**References**

1.	Mwingira F, Nkwengulila G, Schoepflin S, Sumari D, Beck HP, Snounou G, et al. Plasmodium falciparum msp1, msp2 and glurp allele frequency and diversity in sub-Saharan Africa. Malar J. 2011;10.

2.	Broumou I, Plaza DF, Färnert A. Genotyping of Plasmodium falciparum to Assess Clone Composition in Parasite Cultures. Methods in Molecular Biology [Internet]. 2022 [cited 2023 Feb 2];2470:51–68. Available from: https://link.springer.com/protocol/10.1007/978-1-0716-2189-9_6

3.	Liljander A, Wiklund L, Falk N, Kweku M, Mrtensson A, Felger I, et al. Optimization and validation of multi-coloured capillary electrophoresis for genotyping of Plasmodium falciparum merozoite surface proteins (msp1 and 2). Malar J [Internet]. 2009 Apr 23 [cited 2022 Oct 14];8(1):1–14. Available from: https://malariajournal.biomedcentral.com/articles/10.1186/1475-2875-8-78

4.	Sonden K, Doumbo S, Hammar U, Vafa Homann M, Ongoiba A, Traord B, et al. Asymptomatic Multiclonal Plasmodium falciparum Infections Carried Through the Dry Season Predict Protection Against Subsequent Clinical Malaria. Journal of Infectious Diseases. 2015;212(4):608–16.

5.	Kimenyi KM, Wamae K, Ngoi JM, de Laurent ZR, Ndwiga L, Osoti V, et al. Maintenance of high temporal Plasmodium falciparum genetic diversity and complexity of infection in asymptomatic and symptomatic infections in Kilifi, Kenya from 2007 to 2018. Malar J. 2022 Dec 1;21(1).

6.	Schoepflin S, Valsangiacomo F, Lin E, Kiniboro B, Mueller I, Felger I. Comparison of Plasmodium falciparum allelic frequency distribution in different endemic settings by high-resolution genotyping. Malar J [Internet]. 2009 [cited 2025 Jun 12];8(1):250. Available from: https://pmc.ncbi.nlm.nih.gov/articles/PMC2774868/

7.	Rodi M, Kawecka K, Stephan L, Berner L, Medina MS, Lalremruata A, et al. Genetic diversity of Plasmodium malariae in sub-Saharan Africa: a two-marker genotyping approach for molecular epidemiological studies. Front Cell Infect Microbiol. 2024 Jul 19;14:1405198.

8.	Butler JM. The use of capillary electrophoresis in genotyping STR loci. Methods Mol Biol [Internet]. 1998 [cited 2025 Jun 12];98:279–89. Available from: https://link.springer.com/protocol/10.1385/0-89603-443-7:279

9.	Ferreira MU, Hartl DL. Plasmodium falciparum: worldwide sequence diversity and evolution of the malaria vaccine candidate merozoite surface protein-2 (MSP-2). Exp Parasitol [Internet]. 2007 Jan [cited 2024 Oct 28];115(1):32–40. Available from: https://pubmed.ncbi.nlm.nih.gov/16797008/

10.	Cock PJA, Chilton JM, Grüning B, Johnson JE, Soranzo N. NCBI BLAST+ integrated into Galaxy. Gigascience [Internet]. 2015 [cited 2021 Jun 7];4(1). Available from: https://pubmed.ncbi.nlm.nih.gov/26336600/

11.	Camacho C, Coulouris G, Avagyan V, Ma N, Papadopoulos J, Bealer K, et al. BLAST+: Architecture and applications. BMC Bioinformatics [Internet]. 2009 Dec 15 [cited 2021 Jun 7];10. Available from: https://pubmed.ncbi.nlm.nih.gov/20003500/

12.	Altschul SF, Madden TL, Schäffer AA, Zhang J, Zhang Z, Miller W, et al. Gapped BLAST and PSI-BLAST: A new generation of protein database search programs [Internet]. Vol. 25, Nucleic Acids Research. Oxford University Press; 1997 [cited 2021 Jun 7]. p. 3389–402. Available from: https://academic.oup.com/nar/article/25/17/3389/1061651

13.	Guerra CA, Hay SI, Lucioparedes LS, Gikandi PW, Tatem AJ, Noor AM, et al. Assembling a global database of malaria parasite prevalence for the Malaria Atlas Project. Vol. 6, Malaria Journal. 2007.

14.	Hay SI, Guerra CA, Gething PW, Patil AP, Tatem AJ, Noor AM, et al. A world malaria map: plasmodium falciparum endemicity in 2007. PLoS Med. 2009 Mar;6(3):0286–302.

15.	Hadley Wickham. ggplot2: Elegant Graphics for Data Analysis. J R Stat Soc Ser A Stat Soc. 216AD;174(1):245–6.

16.	Kassambara A. “ggplot2” Based Publication Ready Plots [R package ggpubr version 0.6.0]. 2023 Feb 10 [cited 2024 Aug 9]; Available from: https://CRAN.R-project.org/package=ggpubr

17.	Pedersen TL. The Composer of Plots [R package patchwork version 1.2.0]. 2024 Jan 8 [cited 2024 Aug 9]; Available from: https://CRAN.R-project.org/package=patchwork

18.	Wickham H, Pedersen T, Seidel D. scales: Scale Functions for Visualization [Internet]. R package. 2025. Available from: https://scales.r-lib.org/

19.	ggdist: Visualizations of distributions and uncertainty. [cited 2024 Aug 9]; Available from: https://zenodo.org/records/10782896

20.	Kay M. ggdist: Visualizations of Distributions and Uncertainty in the Grammar of Graphics. IEEE Trans Vis Comput Graph. 2024 Jan 1;30(1):414–24.

21.	Oksanen J, Simpson G, Blanchet F, Kindt R, Legendre P, Minchin P, et al. vegan: Community Ecology Package [Internet]. 2025 [cited 2025 Apr 22]. Available from: https://vegandevs.github.io/vegan/.

22.	Aho K. asbio: A Collection of Statistical Tools for Biologists. CRAN: Contributed Packages [Internet]. 2010 Jan 4 [cited 2025 Apr 22]; Available from: https://CRAN.R-project.org/package=asbio

23.	Brunson JC, Read QD. ggalluvial: Alluvial Plots in “ggplot2.” CRAN: Contributed Packages [Internet]. 2017 Nov 26 [cited 2025 Apr 22]; Available from: https://CRAN.R-project.org/package=ggalluvial

24.	Pavoine S. adiv: An r package to analyse biodiversity in ecology. Methods Ecol Evol [Internet]. 2020 Sep 1 [cited 2025 Apr 22];11(9):1106–12. Available from: https://onlinelibrary.wiley.com/doi/full/10.1111/2041-210X.13430

25.	Katoh K, Kuma KI, Toh H, Miyata T. MAFFT version 5: improvement in accuracy of multiple sequence alignment. Nucleic Acids Res [Internet]. 2005 Jan 1 [cited 2024 Oct 16];33(2):511–8. Available from: https://dx.doi.org/10.1093/nar/gki198

26.	Katoh K, Misawa K, Kuma KI, Miyata T. MAFFT: a novel method for rapid multiple sequence alignment based on fast Fourier transform. Nucleic Acids Res [Internet]. 2002 Jul 15 [cited 2024 Oct 16];30(14):3059–66. Available from: https://dx.doi.org/10.1093/nar/gkf436

27.	Katoh K, Standley DM. MAFFT Multiple Sequence Alignment Software Version 7: Improvements in Performance and Usability. Mol Biol Evol [Internet]. 2013 Apr 1 [cited 2024 Oct 16];30(4):772–80. Available from: https://dx.doi.org/10.1093/molbev/mst010

28.	Bahl A, Brunk B, Crabtree J, Fraunholz MJ, Gajria B, Grant GR, et al. PlasmoDB: The Plasmodium genome resource. A database integrating experimental and computational data [Internet]. Vol. 31, Nucleic Acids Research. Nucleic Acids Res; 2003 [cited 2021 May 19]. p. 212–5. Available from: https://pubmed.ncbi.nlm.nih.gov/12519984/

29.	Fraunholz MJ, Roos DS. PlasmoDB: Exploring genomics and post-genomics data of the malaria parasite, Plasmodium falciparum. Redox Report [Internet]. 2003 [cited 2021 May 19];8(5):317–20. Available from: https://pubmed.ncbi.nlm.nih.gov/14962373/

30.	Aurrecoechea C, Brestelli J, Brunk BP, Dommer J, Fischer S, Gajria B, et al. PlasmoDB: A functional genomic database for malaria parasites. Nucleic Acids Res [Internet]. 2009 [cited 2021 May 19];37(SUPPL. 1). Available from: https://pubmed.ncbi.nlm.nih.gov/18957442/

31.	Bahl A, Brunk B, Coppel RL, Crabtree J, Diskin SJ, Fraunholz MJ, et al. PlasmoDB: The Plasmodium genome resource. An integrated database providing tools for accessing, analyzing and mapping expression and sequence data (both finished and unfinished). Nucleic Acids Res [Internet]. 2002 Jan 1 [cited 2021 May 19];30(1):87–90. Available from: https://pubmed.ncbi.nlm.nih.gov/11752262/

32.	Minh BQ, Nguyen MAT, Von Haeseler A. Ultrafast Approximation for Phylogenetic Bootstrap. Mol Biol Evol [Internet]. 2013 May 1 [cited 2025 Mar 10];30(5):1188–95. Available from: https://dx.doi.org/10.1093/molbev/mst024

33.	Kalyaanamoorthy S, Minh BQ, Wong TKF, Von Haeseler A, Jermiin LS. ModelFinder: Fast model selection for accurate phylogenetic estimates. Nat Methods [Internet]. 2017 May 30 [cited 2021 May 20];14(6):587–9. Available from: https://pubmed.ncbi.nlm.nih.gov/28481363/

34.	Schrempf D, Minh BQ, De Maio N, von Haeseler A, Kosiol C. Reversible polymorphism-aware phylogenetic models and their application to tree inference. J Theor Biol. 2016 Oct 21;407:362–70.

35.	Nguyen LT, Schmidt HA, Von Haeseler A, Minh BQ. IQ-TREE: A Fast and Effective Stochastic Algorithm for Estimating Maximum-Likelihood Phylogenies. Mol Biol Evol [Internet]. 2015 Jan 1 [cited 2025 Mar 10];32(1):268–74. Available from: https://dx.doi.org/10.1093/molbev/msu300

36.	Hoang DT, Chernomor O, Von Haeseler A, Minh BQ, Vinh LS. UFBoot2: Improving the Ultrafast Bootstrap Approximation. Mol Biol Evol [Internet]. 2018 Feb 1 [cited 2025 Mar 10];35(2):518–22. Available from: https://dx.doi.org/10.1093/molbev/msx281

37.	Guindon S, Dufayard JF, Lefort V, Anisimova M, Hordijk W, Gascuel O. New Algorithms and Methods to Estimate Maximum-Likelihood Phylogenies: Assessing the Performance of PhyML 3.0. Syst Biol [Internet]. 2010 May 1 [cited 2025 Mar 10];59(3):307–21. Available from: https://dx.doi.org/10.1093/sysbio/syq010

38.	Letunic I, Bork P. Interactive Tree of Life (iTOL) v4: Recent updates and new developments. Nucleic Acids Res [Internet]. 2019 Jul 1 [cited 2021 May 28];47(W1):W256–9. Available from: https://academic.oup.com/nar/article/47/W1/W256/5424068

39.	Letunic I, Bork P. Interactive Tree of Life (iTOL) v6: recent updates to the phylogenetic tree display and annotation tool. Nucleic Acids Res [Internet]. 2024 Jul 5 [cited 2025 Feb 12];52(W1):W78–82. Available from: https://dx.doi.org/10.1093/nar/gkae268
